# Canonical WNT Ligands Produced by Regulatory T Cells Restrain Effector CD4^+^ T Cell Responses

**DOI:** 10.64898/2026.04.12.717050

**Authors:** Kashish Singh Parihar, Margaret R. Niemeier, Igal Ifergan

**Affiliations:** Department of Immunology, Cincinnati Children’s Hospital Medical Center, University of Cincinnati College of Medicine, Cincinnati, OH USA; Department of Molecular and Cellular Biosciences, College of Medicine, University of Cincinnati, Cincinnati, OH USA; Neuroscience Graduate Program, College of Medicine, University of Cincinnati, Cincinnati, OH USA

**Keywords:** WNT, FZD, Tregs, CD4^+^ T cells, immune suppression

## Abstract

Regulatory T cells (Tregs) are essential for maintaining immune homeostasis by suppressing excessive activation of effector T cells. Although several mechanisms of Treg-mediated suppression have been described, the molecular signals that contribute to this regulation remain incompletely understood. WNT signaling, best known for its roles in development and tissue homeostasis, has recently emerged as an important regulator of immune function, but its contribution to Treg-mediated immune suppression is largely unknown.

Here, we show that Tregs preferentially express multiple canonical WNT ligands, including WNT2B, WNT3, WNT7B, and WNT10B, compared with conventional CD4^+^ T cells. These WNT proteins were detected intracellularly in Tregs, and WNT2B and WNT3 were actively secreted into culture supernatants. Conventional CD4^+^ T cells expressed Frizzled receptors capable of sensing these ligands. Pharmacological inhibition of canonical WNT signaling using the antagonist mDKK-1 enhanced CD4^+^ T cell activation and proliferation and increased pro-inflammatory cytokine expression, while anti-inflammatory IL-10 remained unchanged.

Together, these findings identify Tregs as a source of canonical WNT ligands and suggest that Treg-derived WNT signaling contributes to the suppression of effector CD4^+^ T cell responses. This work reveals a previously underappreciated pathway through which Tregs regulate immune activity and identifies WNT signaling as a potential target for modulating inflammatory immune responses.

## 1. Introduction

The immune system relies on tightly regulated cellular interactions to mount effective responses against pathogens while preserving tolerance to self. Within the adaptive immune system, CD4^+^ T helper cells play a central role in shaping immune outcomes through their differentiation into specialized functional subsets. Among these, regulatory T cells (Tregs) are essential for maintaining immune homeostasis by limiting excessive or inappropriate immune activation^1^.

Tregs are defined by the expression of CD4, high levels of CD25, and the transcription factor FOXP3 (Forkhead Box P3)^2^, and they suppress immune responses through multiple mechanisms. These include modulation of antigen-presenting cell function via inhibitory molecules such as CTLA-4 (Cytotoxic T-Lymphocyte Associated protein 4)^3^ and LAG-3 (Lymphocyte Activation Gene-3)^4^, metabolic competition for IL-2 availability^5^, and secretion of anti-inflammatory cytokines including IL-10^6–8^, TGF-β^9, 10^, and IL-35^11, 12^. Through these pathways, Tregs restrain effector T cell expansion, prevent immunopathology, and preserve peripheral tolerance. Although these canonical suppressive mechanisms have been extensively characterized, additional signaling pathways likely contribute to Treg biology and function.

One such pathway is WNT signaling, a conserved network of secreted glycoproteins best known for its roles in embryonic development, tissue homeostasis, and stem cell maintenance. The WNT family comprises 19 ligands that signal through Frizzled receptors and co-receptors to activate either β-catenin-dependent (canonical) or β-catenin-independent (non-canonical) pathways. Canonical WNT signaling regulates transcriptional programs involved in proliferation, survival, and differentiation, whereas non-canonical pathways influence cytoskeletal organization, migration, and calcium-dependent signaling. Beyond development, accumulating evidence indicates that WNT signaling also contributes to immune regulation.

Recent studies have shown that immune cells can both respond to and produce WNT ligands, thereby establishing potential autocrine and paracrine regulatory circuits. For instance, macrophages upregulate WNT3A^13^ and WNT7B^14^ following tissue injury, where these ligands facilitate regeneration and repair. Activated T cells have been shown to secrete WNT10B^15^ under specific co-stimulatory conditions, thereby influencing tissue repair and shaping effector responses. Dendritic cells express WNT3A, and activation of β-catenin signaling in dendritic cells promotes the expression of tolerogenic mediators that support regulatory T cell generation^16, 17^. B lymphocytes and their progenitors respond to WNT3A by stabilizing β-catenin and express multiple WNT ligands (WNT2B, 5B, 8A, 10A, 16), positioning them as active sources of canonical signals within lymphoid tissues^18^. Additionally, monocytes have been shown to secrete WNT3A in inflammatory contexts, thereby enhancing β-catenin signaling in surrounding immune and stromal cells^19^. Meanwhile, neutrophils upregulate WNT5A during infection and tissue injury, contributing to tissue remodeling and immune regulation^20^. Collectively, these findings indicate that immune cells are not only targets of WNT signals but can also serve as active producers, further linking WNT signaling to immune regulation.

Despite these advances, the expression and functional contribution of WNT ligands within CD4^+^ T cell subsets, particularly Tregs, remain poorly defined. Here, we examined whether Tregs express and secrete specific WNT ligands that contribute to their suppressive function by systematically assessing WNT gene and protein expression and evaluating the impact of Treg-derived WNT signals on conventional CD4^+^ T cell activation.

## 2. Materials and Methods

### 2.1 Mice

Male and female C57BL/6 mice (Jackson Laboratory, #000664), aged 7-12 weeks, were maintained under specific pathogen-free conditions at the University of Cincinnati’s Laboratory Animal Medical Services (LAMS). All procedures were approved by the Institutional Animal Care and Use Committee (IACUC) and performed in accordance with AAALAC guidelines.

### 2.2 In vitro experiments

Spleens were harvested from mice and physically homogenized through a 100 µm cell strainer (Corning Inc.) to obtain single-cell suspensions, followed by red blood cell lysis with 0.83% ammonium chloride. CD4⁺ T cells were isolated using either the EasySep Mouse CD4⁺ T Cell Negative Isolation Kit (StemCell Technologies; cat# 19852RF) or the MojoSort CD4⁺ T Cell Negative Isolation Kit (BioLegend; cat# 480005), following the manufacturers’ protocols, and further enriched for naïve CD4⁺ T cells via positive selection with CD62L microbeads (Miltenyi Biotec; cat# 130-049-701).

For *in vitro* differentiation, naïve CD4^+^ T cells were plated at 0.2 - 0.5 × 10^6^ cells per well in RPMI-1640 medium supplemented with 10% FBS, 2 mM L-glutamine (Gibco), and 100 U/mL penicillin-streptomycin (Gibco), and stimulated with plate-bound anti-CD3 (2.5 µg/mL; BD Biosciences) and soluble anti-CD28 (10 µg/mL; BioLegend) antibodies.

Th0 differentiation was induced with 200 U/mL recombinant mouse IL-2 (rmIL-2, R&D Systems). Th1, Th17, and Treg polarizations were achieved by adding cytokine and antibody combinations as follows: Th1: 200 U/mL rmIL-2 (R&D Systems), 10 ng/mL rmIL-12 (PeproTech), 1 µg/mL anti-IL-4 (BioLegend); Th17: 1 ng/mL TGF-β1 (PeproTech), 20 ng/mL IL-6 (PeproTech), 20 ng/mL IL-23 (PeproTech), 20 ng/mL IL-1β (PeproTech) 1 µg/mL anti-IFN-γ (BioLegend), 1 µg/mL anti-IL-4 (BioLegend); Treg: 200 U/mL IL-2 (R&D Systems), 26 ng/mL TGF-β1 (PeproTech), 1 µg/mL anti-IFN-γ (BioLegend). Cells were incubated for 96 hours at 37°C with 5% CO₂ to support differentiation.

Where applicable, cultures were treated with the WNT antagonist mDKK-1 (250 ng/mL; R&D Systems).

### 2.3 FZD Receptor Expression

Purified CD4⁺ T cells were cultured under resting conditions or stimulated with plate-bound anti-CD3 (2.5 µg/mL) and soluble anti-CD28 (10 µg/mL). Resting cells were harvested after 24 hours, whereas activated cells were harvested after 48 hours. Cells were stained with antibodies specific for FZD1 - FZD10 receptors as follows:

- FZD-1: APC-conjugated, clone 162531 (R&D Systems; cat# FAB11201A); isotype Rat IgG2a
- FZD-2: Rabbit polyclonal (Novus Biologicals; cat# NLS3488); isotype Rabbit IgG
- FZD-3: APC-conjugated, clone 169310 (R&D Systems; cat# FAB1001A); isotype Rat IgG2a
- FZD-4: Alexa647-conjugated, clone 145901 (R&D Systems; cat# FAB194R); isotype Rat IgG2a
- FZD-5: Rabbit polyclonal (Sigma; cat# SAB4503132); isotype Rabbit IgG
- FZD-6: Goat polyclonal (R&D Systems; cat# AF1526); isotype Goat IgG
- FZD-7: APC-conjugated, clone 151143 (R&D Systems; cat# FAB1981A); isotype Rat IgG2a
- FZD-8: Rabbit polyclonal (Novus Biologicals; cat# NLS4767); isotype Rabbit IgG
- FZD-9: Rabbit polyclonal (Novus Biologicals; cat# NB100-59000); isotype Rabbit IgG
- FZD-10: Rabbit polyclonal (ThermoFisher; cat# PA5-75325); isotype Rabbit IgG For directly conjugated antibodies (FZD-1, FZD-3, FZD-4, FZD-7), staining was performed according to the manufacturer’s instructions.

For unconjugated primary antibodies, cells were incubated with the primary antibody at 10 µg/mL for 20 minutes, washed, and then stained with the appropriate fluorochrome-conjugated secondary antibody.

- For rabbit-derived antibodies (FZD-2, FZD-5, FZD-8, FZD-9, FZD-10): goat anti-rabbit IgG APC-conjugated (R&D Systems; cat# F0111)
- For goat-derived antibody (FZD-6): donkey anti-goat IgG (H+L) APC-conjugated (R&D Systems; cat# F0108)

All secondary antibody staining was performed following the manufacturer’s protocols. Corresponding isotype controls were included where appropriate. Flow cytometry was conducted on a BD LSRFortessa system, and data were analyzed using FlowJo software. Expression levels of each FZD receptor were compared between resting and activated CD4⁺ T cells to identify activation-dependent changes in WNT receptor expression.

### 2.4 CFSE Labeling

CFSE labeling was performed using the CellTrace™ CFSE Cell Proliferation Kit (Thermo Fisher; cat# C34554) according to the manufacturer’s instructions, with minor modifications. Briefly, CD4^+^ T cells were labeled at a final dilution of 1:2,000 and incubated at 37 °C for 10 minutes, followed by quenching with FBS and washing prior to culture.

### 2.5 Suppression Assays

CFSE-labeled CD4⁺ T cells were stimulated with plate-bound anti-CD3 (2.5 µg/mL) and soluble anti-CD28 (10 µg/mL) antibodies and cultured either alone, co-cultured with Th0 or Treg cells (1:1 ratio), or exposed to conditioned supernatants collected from differentiated T cell cultures. Where indicated, mDKK-1 (250ng/mL) was added.

### 2.6 Quantitative Real-Time PCR

Cells were lysed with TRIzol Reagent (Ambion), and total RNA was extracted following the Invitrogen TRIzol LS manufacturer’s protocol. An additional phenol-based cleanup was performed to improve RNA purity. RNA quality was assessed using a NanoDrop spectrophotometer (260/280 nm), and only samples with purity ratios between 1.7 and 1.9 were used for cDNA synthesis. Complementary DNA (cDNA) was synthesized from 300 ng of total RNA per sample with volumes normalized based on mRNA concentration, using the RT² First Strand Kit (Qiagen) according to the manufacturer’s instructions.

Mouse-specific RT² PCR arrays (cat# 330171, CLAM42845, SABiosciences, Qiagen) were employed to profile 19 WNT genes. Arrays included five housekeeping genes, TGF-β, genomic DNA, reverse transcription efficiency controls, and a positive PCR control.

Quantitative PCR was performed on a QuantStudio 3 system (Thermo Fisher) with the following cycling conditions: 95 °C for 10 min, followed by 40 cycles of 95 °C for 15 s and 60 °C for 1 min, with fluorescence recorded during each cycle. Data were analyzed using Qiagen’s GeneGlobe Design and Analysis Tool. Ct values above 35 were excluded, and B2m and β-actin were used for normalization. Fold changes in gene expression were calculated relative to naïve CD4⁺ T cells, with ±1.5-fold used as a cutoff for biologically relevant changes.

Genes of interest were further validated across naïve CD4⁺ T cells, Tregs, Th0, Th1, and Th17 cells using custom primers (Integrated DNA Technologies, IDT) in 96-well MicroAmp Optical Plates (Thermo Fisher). Primer sequences were:

- WNT2B: F 5’-CCA TTA CGG TGT TCG CTT TGC C-3’; R 5’-CAG CTT CAG GAA TCT CCG AAC AG-3’
- WNT3: F 5’-CAA GCA CAA CAA TGA AGC AGG C-3’; R 5’-TCG GGA CTC ACG GTG TTT CTC-3’
- WNT7B: F 5’-TTC TCG TCG CTT TGT GGA TGC C-3’; R 5’-CAC CGT GAC ACT TAC ATT CCA GC-3’
- WNT9B: F 5’-AAG TAC AGC ACC AAG TTC CTC AGC-3’; R 5’-GAA CAG CAC AGG AGC CTG ACA C-3’
- WNT10B: F 5’-ACC ACG ACA TGG ACT TCG GAG A-3’; R 5’-CCG CTT CAG GTT TTC CGT TAC C-3’
- β-Actin: F 5’-GGC TGT ATT CCC CTC CAT CG-3’; R 5’-CCA GTT GGT AAC AAT GCC ATG T-3’

Primer mixes were prepared by combining 1 µL each of 10 µM forward and reverse primers with 8 µL nuclease-free water. Each qPCR reaction contained 1 µL of cDNA (synthesized from a total of 350 ng mRNA per sample), 1 µL primer mix, 5 µL SYBR Green (Thermo Fisher), and 3 µL nuclease-free water, totaling 10 µL per well. qPCR was performed on the QuantStudio 3 with the following program: 50 °C for 2 min, 95 °C for 2 min, and 40 cycles of 95 °C for 15 s and 60 °C for 1 min, followed by melt curve analysis (95 °C for 15 s, 60 °C for 1 min, 95 °C for 15 s). Relative gene expression was calculated using the ΔΔCt method, normalized to β-actin, and expressed relative to naïve CD4⁺ T cells.

### 2.7 Flow Cytometry

For analysis of cytokine production, T cells were stimulated with phorbol 12-myristate 13-acetate (PMA; 20ng/mL) and ionomycin (1 µg/mL) for 4 hours. Brefeldin A (2 µg/mL; Sigma) was added during the final 2 hours of stimulation to inhibit cytokine secretion.

Cells were stained with LIVE/DEAD Fixable Aqua dye (Thermo Fisher; cat# L34957) for 20 minutes at room temperature in the dark. Surface staining was then performed using fluorochrome-conjugated antibodies (BioLegend or Thermo Fisher) for 20 minutes at 4 °C in FACS buffer (PBS supplemented with 1% FBS and 0.1% sodium azide).

For intracellular cytokine staining, cells were fixed and permeabilized using BD Cytofix/Cytoperm reagents (BD Biosciences) according to the manufacturer’s instructions. FOXP3 staining was performed using the eBioscience Foxp3/Transcription Factor Staining Buffer Set.

To detect intracellular WNT proteins, cells were stained with primary antibodies against WNT2B (RRID: AB_2911621), WNT3 (RRID: AB_2900762), WNT7B (RRID: AB_2852816), WNT9B (RRID: AB_2900764), and WNT10B (RRID: AB_2718315) (Thermo Fisher) each at a final concentration of 10 µg/mL. A rabbit IgG isotype control antibody (R&D Systems) was included to assess non-specific binding. After incubation with primary WNT antibodies, cells were stained with APC-conjugated goat anti-rabbit IgG secondary antibody (R&D Systems) in 1X permeabilization buffer.

Data were acquired on a BD FACSCanto II and analyzed using FlowJo software (v10.10).

### 2.8 ELISA

WNT secretion was measured in culture supernatants, using ELISA kits for WNT2B (RayBiotech; cat# ELM-WNT2B), and WNT3 (CUSABIO; cat# CSB-EL026135M0) according to the manufacturer’s protocols. Attempts to quantify WNT7B and WNT10B using available commercial ELISA kits did not yield detectable signals.

Cytokine levels (TNF-α, GM-CSF, IFN-γ, IL-10) were quantified by ELISA using commercially available kits (BioLegend) according to the manufacturer’s instructions.

Absorbance at 450 nm was recorded using a Biotek Epoch microplate reader, and cytokine concentrations were calculated from standard curves.

### 2.9 Statistical Analysis

Statistical analyses were performed using GraphPad Prism version 10.0 (GraphPad Software). Comparisons between groups were conducted using paired or unpaired t-tests, as well as two-way ANOVA as appropriate. Differences were considered statistically significant only when p-values were below 0.05.

## 3. Results

### 3.1 Tregs preferentially express canonical WNT ligands

To determine whether Tregs express WNT ligands, we first isolated CD62L^+^ CD44^lo^ naïve CD4^+^ T cells from healthy adult mice (**Fig. S1**) and differentiated them into Tregs. Cell identity and purity were confirmed by flow cytometry, and only populations with ≥80% purity were used for subsequent analyses (**Fig. S2**).

We first compared WNT gene expression in Tregs and naïve CD4^+^ T cells by qPCR, normalizing expression to the housekeeping genes *B2m* and *Actb*. Tregs displayed a distinct WNT expression profile relative to naïve CD4^+^ T cells, as summarized in the heatmap shown in **Fig. S3**, and quantitative fold changes with associated p-values are provided in **Table 1**. Among the 19 WNT genes examined, *WNT2B*, *WNT3*, and *WNT10B* were significantly upregulated in Tregs compared with naïve CD4^+^ T cells (*p* < 0.05), indicating preferential enrichment within the regulatory lineage. *WNT7B* displayed a trend toward increased expression (*p* = 0.07) and was included in subsequent analyses based on its established role in immune signaling. In contrast, *WNT9B*, another canonical WNT family member, did not reach statistical significance.

As an internal validation of Treg identity, *Tgfb1*, a well-characterized marker of Treg function, was significantly elevated, confirming that the transcriptional profiles captured key regulatory features. Although *WNT11* expression was increased, it was excluded from further analysis because it primarily participates in non-canonical WNT signaling pathways^21, 22^, which were outside the scope of this study. Notably, the remaining upregulated WNT genes^23–28^ are predominantly associated with β-catenin-dependent signaling, consistent with enrichment of canonical WNT pathway components in Tregs.

To extend these findings, we examined the expression of selected WNT genes across multiple CD4^+^ T cell subsets. Transcript levels of *WNT2B*, *WNT3*, *WNT7B*, *WNT9B* and *WNT10B* were quantified by qPCR in naïve CD4^+^ T cells, Th0, Th1, Th17, and Treg populations (**Fig. 1**). Consistent with the initial screen, *WNT2B*, *WNT3*, *WNT7B*, and *WNT10B* were all elevated in Tregs relative to naïve CD4^+^ T cells. *WNT7B* and *WNT10B* showed the strongest Treg-selective pattern, as their expression was significantly higher in Tregs than in the other CD4^+^ T cell subsets examined. *WNT2B* and *WNT3* were also elevated in Th0 cells, indicating that these ligands are upregulated in Tregs but are not uniquely restricted to the regulatory lineage. *WNT9B* did not show a distinct subset-specific pattern. Together, these data indicate that Tregs are enriched for canonical WNT ligand expression, with WNT7B and WNT10B showing the clearest Treg-associated enrichment.

**Figure 1.**
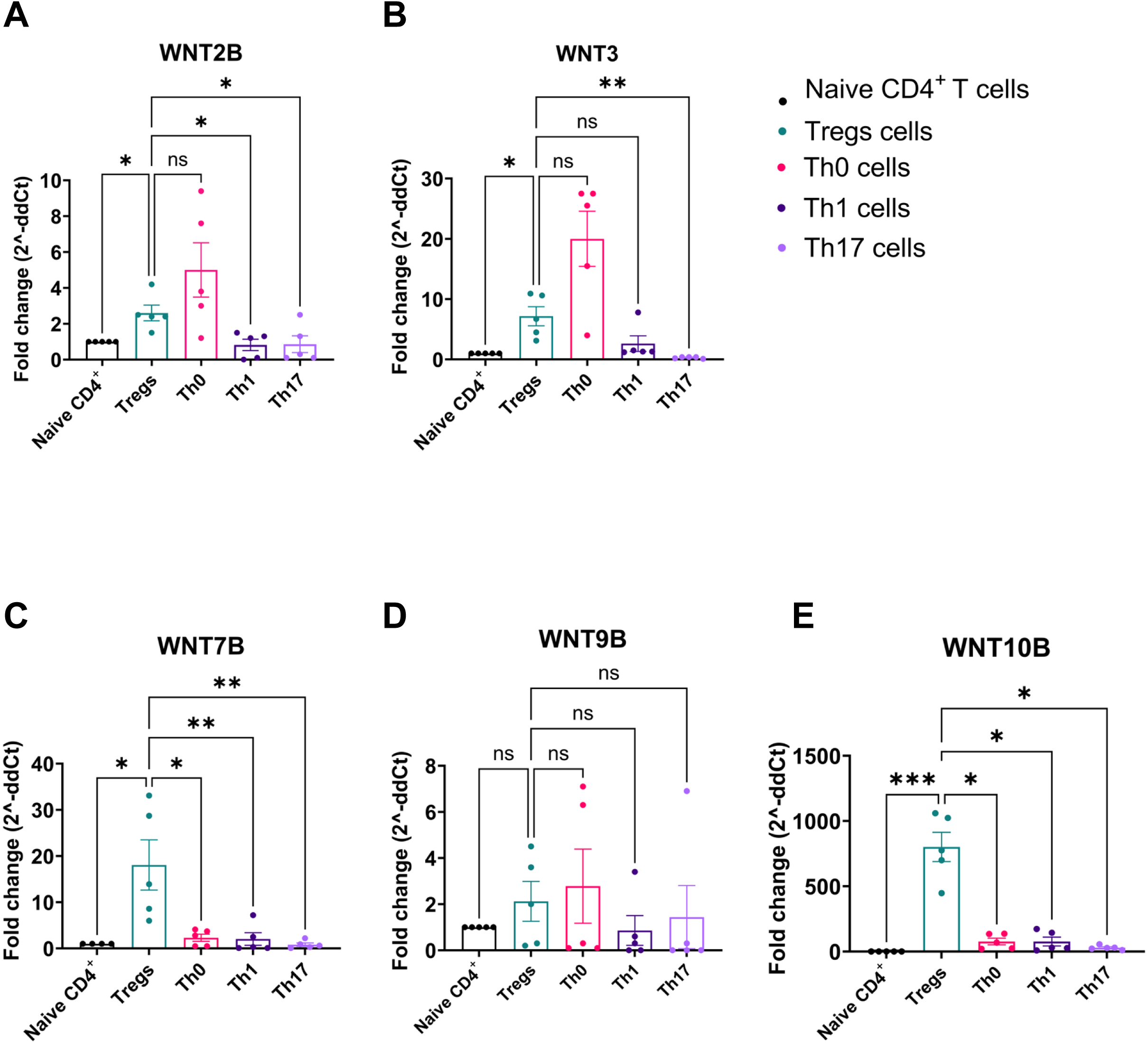
Differential WNT gene expression across CD4^+^ T cell subsets. Relative expression of (**A**) *WNT2B*, (**B**) *WNT3*, (**C**) *WNT7B*, (**D**) *WNT9B*, and (**E**) *WNT10B* was quantified by qPCR in CD4^+^ T cell subsets and expressed as fold change (2^-ΔΔCt). Naïve CD4^+^ T cells are shown in black, regulatory T cells (Tregs) in green, Th0 cells in pink, Th1 cells in violet, and Th17 cells in lilac. Data represent mean ± SEM from five independent experiments (n = 5). Statistical significance was assessed using two-way ANOVA (\**p* < 0.05, \*\**p* < 0.01).

We next asked whether these transcriptional differences were reflected at the protein level. Intracellular flow cytometric analysis of naïve CD4^+^ T cells and *ex vivo* natural Tregs (nTregs), using the gating strategy shown in **Fig. S4**, revealed differential expression of WNT proteins. Representative histogram overlays are shown in **Fig. 2A**. Quantification demonstrated significantly higher frequencies of WNT2B, WNT3, WNT7B, and WNT10B expression in nTregs compared with naïve CD4^+^ T cells **(Fig. 2B-D, F)**, while WNT9B expression remained unchanged **(Fig. 2E)**. These findings confirm that transcriptional enrichment of canonical WNT ligands in Tregs is reflected at the protein level.

**Figure 2.**
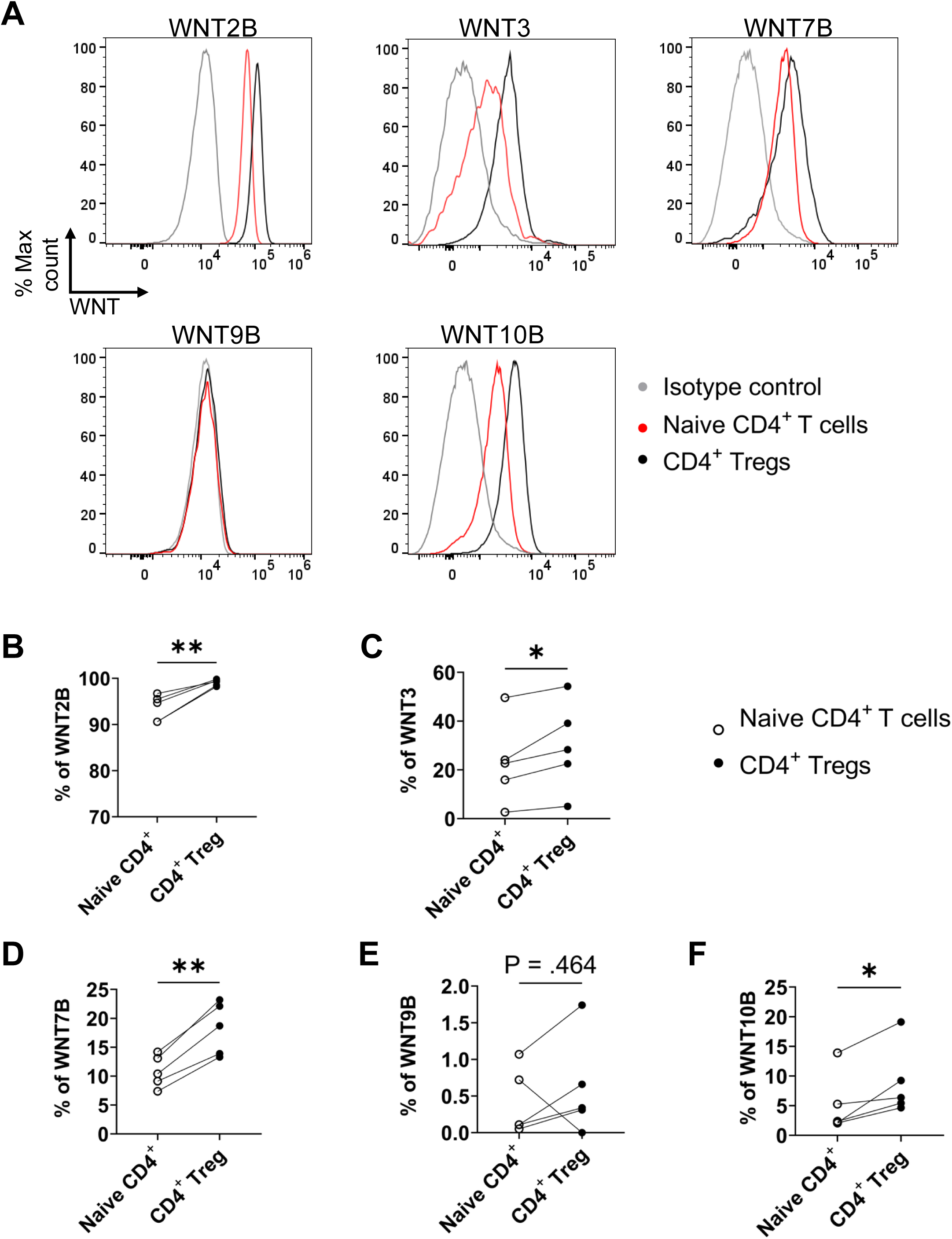
Characterization of WNT protein expression in naïve CD4^+^ T cells and regulatory T cells. (**A**) Representative histogram overlays of intracellular WNT protein expression in naïve CD4^+^ T cells and natural regulatory T cells (nTregs). Isotype controls are shown in grey, naïve CD4^+^ T cells in red, and nTregs in black. (**B-F**) Quantification of the percentage of cells expressing (**B**) WNT2B, (**C**) WNT3, (**D**) WNT7B, (**E**) WNT9B, and (**F**) WNT10B. Data are presented as mean ± SEM from five independent experiments (n = 5). Statistical significance was determined using paired t-tests (\**p* < 0.05, \*\**p* < 0.01).

### 3.2 Tregs secrete canonical WNT proteins

We next examined whether differentiated T cells secrete WNT proteins. Because naïve CD4^+^ T cells exhibit limited survival in prolonged culture, we compared supernatants from differentiated Th0 and Treg cultures after confirming their phenotype by flow cytometry (**Fig. S2 and S5)**.

ELISA-based quantification showed that WNT2B levels were significantly higher in Treg supernatants than in Th0 supernatant (**Fig. 3A**). WNT3 secretion was similarly elevated in Treg cultures relative to Th0 controls (**Fig. 3B**). Although *WNT2B* and *WNT3* transcripts were also detected in Th0 cells, ELISA revealed greater accumulation of these proteins in Treg-conditioned supernatants, indicating enhanced secretion by Tregs relative to Th0 cells. In contrast, commercially available ELISA assays for WNT7B and WNT10B did not produce reliable signals despite repeated attempts, precluding confident quantification of these ligands in conditioned media. Together, these results indicate that Tregs preferentially secrete specific canonical WNT ligands, particularly WNT2B and WNT3.

**Figure 3.**
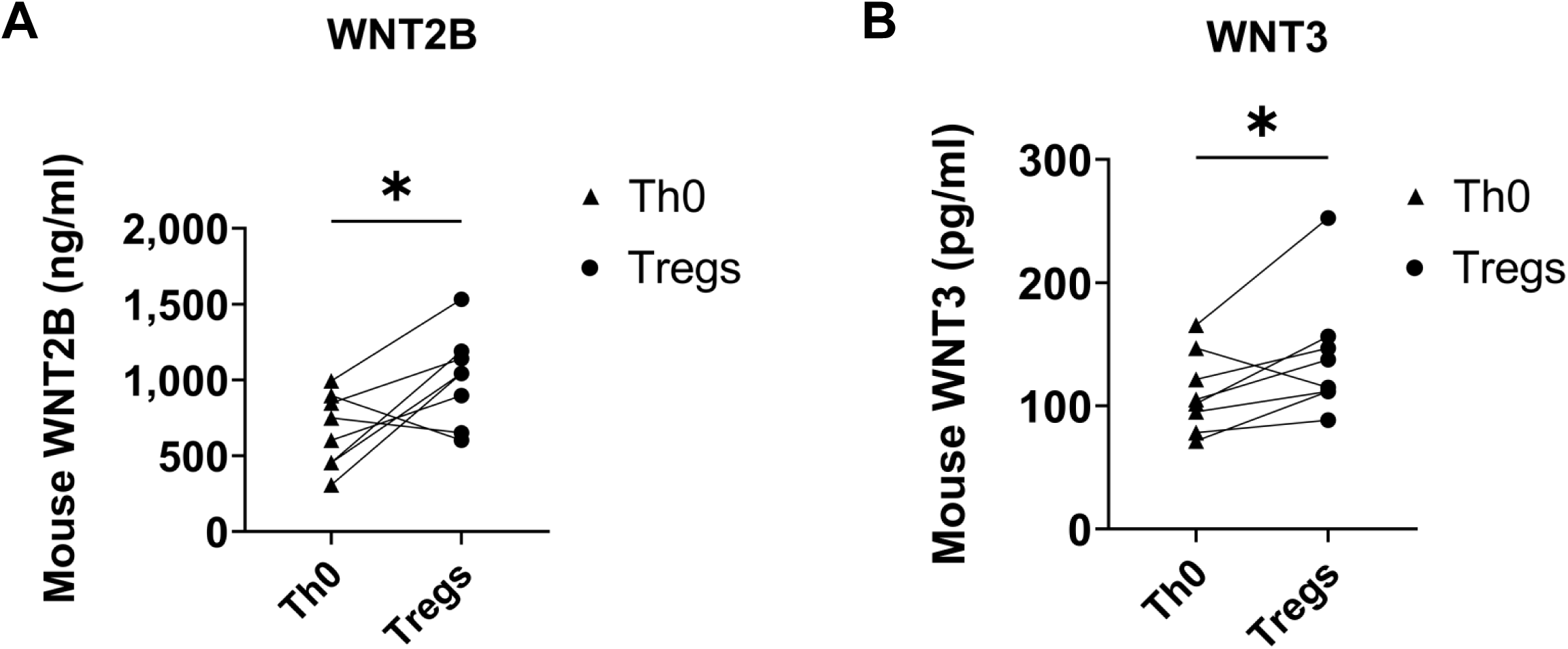
Differential secretion of WNT proteins by Th0 cells and regulatory T cells. Culture supernatants from differentiated Th0 and Treg cells were analyzed by ELISA to quantify secreted WNT proteins. Concentrations of (**A**) WNT2B and (**B**) WNT3 are shown. Each point represents paired samples from independent experiments (n = 8). Data are presented as mean ± SEM. Statistical significance was determined using paired t-tests (\**p* < 0.05).

### 3.3 CD4^+^ T cells express a subset of Frizzled receptors capable of sensing WNT ligands

Having established that Tregs produce canonical WNT ligands, we next asked whether conventional CD4^+^ T cells express receptors capable of receiving these signals. Flow cytometric screening of FZD1-FZD10 revealed that CD4^+^ T cells express a restricted set of Frizzled receptors, with FZD4, FZD5, and FZD10 showing the most prominent expression **(Fig. 4A)**. Upon activation with anti-CD3 and anti-CD28 antibodies, FZD5 and FZD10 increased significantly, while FZD4 was reduced relative to resting cells. Representative histogram overlays for these receptors are shown in **Fig. 4B**. The remaining FZD family members were expressed at low levels or were not significantly altered by activation. These findings indicate that both resting and activated CD4^+^ T cells express FZD receptors capable of sensing WNT ligands in their environment.

**Figure 4.**
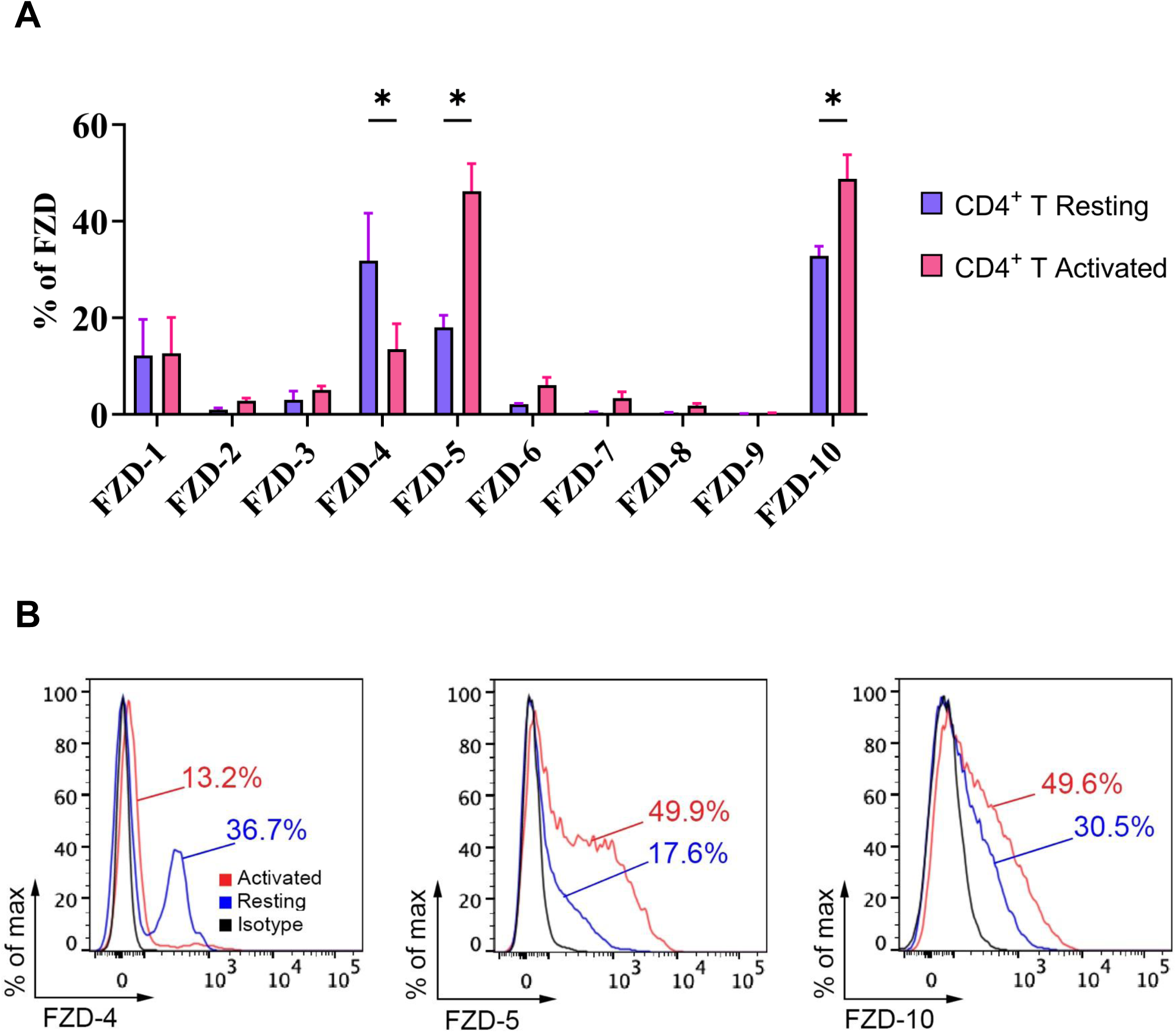
Expression profile of Frizzled (FZD) receptors on CD4^+^ T cells. **(A)** Flow cytometric analysis of surface expression of FZD1-FZD10 on resting and activated CD4^+^ T cells. Expression was quantified as the percentage of FZD-positive CD4^+^ T cells under resting conditions (violet) or following 48 h stimulation with anti-CD3 and anti-CD28 antibodies (pink). Data represent mean ± SEM from five independent experiments (n = 5). Statistical significance was determined using paired t-tests (\**p* < 0.05). **(B)** Representative histogram overlays showing surface expression of FZD4, FZD5, and FZD10 on CD4^+^ T cells. Resting CD4^+^ T cells are shown in blue, activated CD4^+^ T cells in red, and isotype controls in black.

### 3.4 Treg-derived WNT signaling suppresses CD4^+^ T cell activation, proliferation, and inflammatory cytokine production

We next tested whether WNT-dependent signals contribute to Treg-mediated regulation of CD4^+^ T-cell responses. CD4^+^ T cells were cultured alone or co-cultured with *in vitro*-differentiated Tregs at a 1:1 ratio and stimulated with anti-CD3 and anti-CD28 antibodies for 72 hours in the presence or absence of the canonical WNT antagonist mDKK-1, which blocks WNT interaction with LRP5/6 co-receptors^29^. For co-culture experiments, the analyzed responder population was defined as CD4^+^FOXP3^-^ cells to exclude Tregs from the downstream readout (gating strategy in **Fig. S6**). Activation markers and cytokine expression were then assessed in this population (representative flow plots in **Fig. S7**). For clarity, CD4^+^ T cells cultured alone are shown in **Fig. 5**, whereas co-culture conditions are presented in **Fig. 6**.

**Figure 5.**
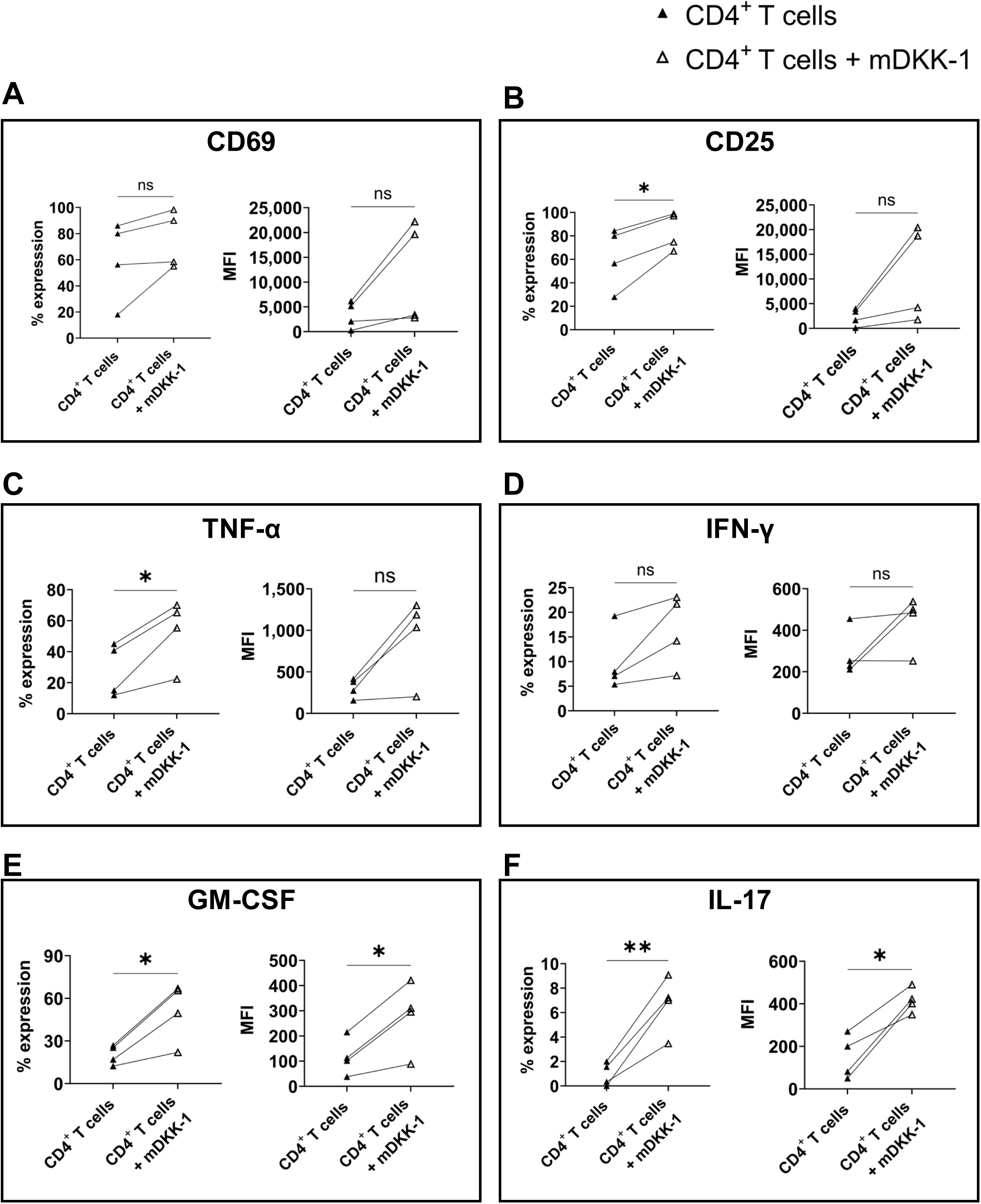
Canonical WNT blockade enhances activation and pro-inflammatory cytokine expression in CD4^+^ T cells cultured alone. CD4^+^ T cells were cultured alone and stimulated with anti-CD3 and anti-CD28 antibodies for 72 h in the presence or absence of the WNT antagonist mDKK-1. Panels show the percentage of positive cells and mean fluorescence intensity (MFI) for activation markers (**A**) CD69 and (**B**) CD25, and cytokines (**C**) TNF-α, (**D**) IFN-γ, (**E**) GM-CSF, and (**F**) IL-17. Data represent mean ± SEM from four independent experiments (n = 4). Statistical significance was determined using paired t-tests (\**p* < 0.05, \*\**p* < 0.01).

**Figure 6.**
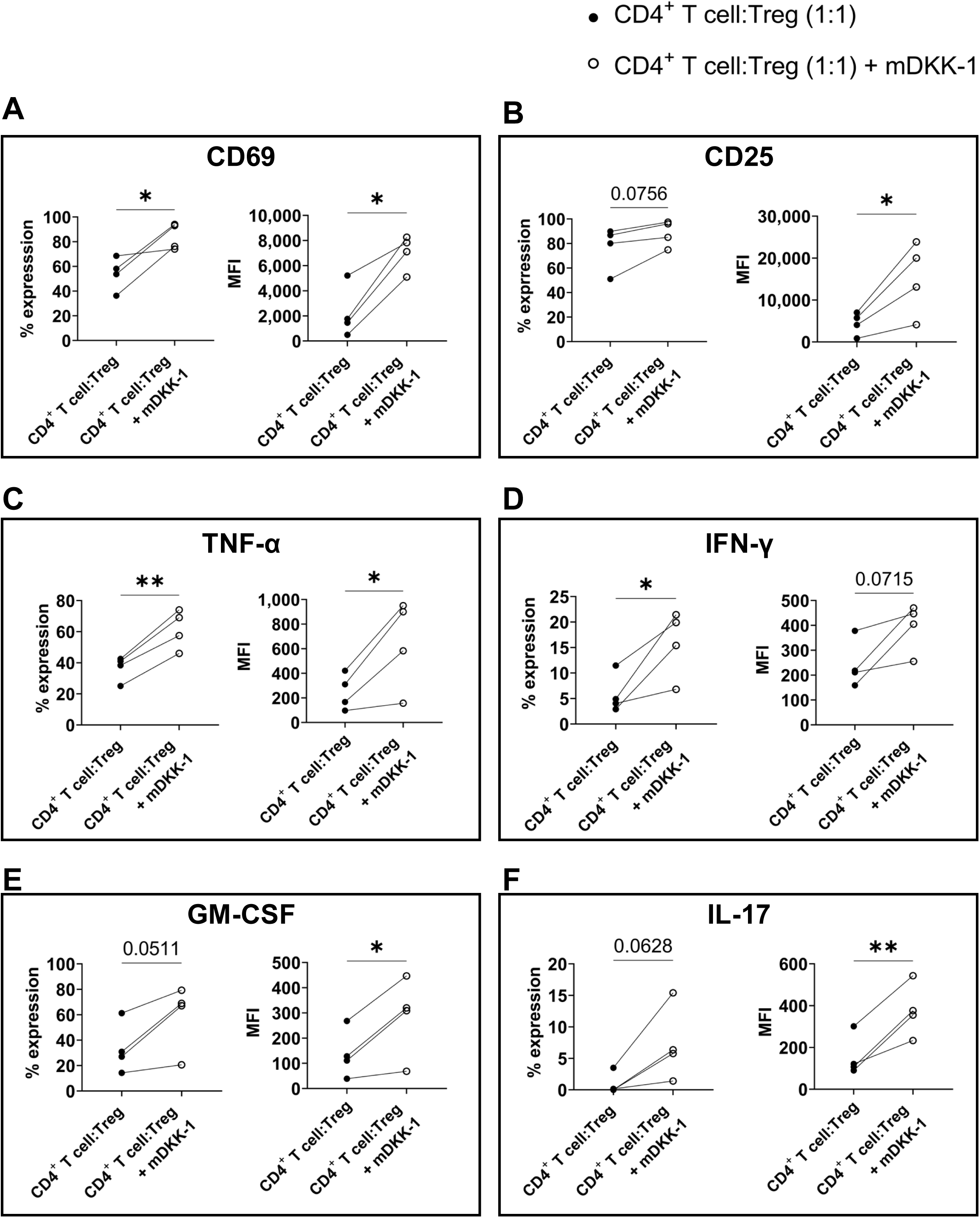
Canonical WNT blockade partially reverses suppression of CD4^+^ T cells co-cultured with Tregs. CD4^+^ T cells were co-cultured with Tregs at a 1:1 ratio and stimulated with anti-CD3 and anti-CD28 antibodies for 72 h in the presence or absence of mDKK-1. Panels show the percentage of positive cells and mean fluorescence intensity (MFI) for activation markers (**A**) CD69 and (**B**) CD25, and cytokines (**C**) TNF-α, (**D**) IFN-γ, (**E**) GM-CSF, and (**F**) IL-17. Data represent mean ± SEM from four independent experiments (n = 4). Statistical significance was determined using paired t-tests (\**p* < 0.05, \*\**p* < 0.01).

When cultured alone, CD4^+^ T cells displayed modest changes in activation following WNT inhibition. CD69 showed a small, non-significant increase, whereas the frequency of CD25^+^ cells increased significantly **(Fig. 5A-B)**. WNT blockade also increased the expression of the pro-inflammatory cytokines TNF-α, GM-CSF, and IL-17, but not IFN-γ **(Fig. 5C-F)**.

Under co-culture conditions, inhibition of WNT signaling with mDKK-1 partially reversed the restrained activation state of CD4^+^ T cells. CD69 expression increased significantly in both frequency and MFI, and CD25 MFI was also elevated **(Fig. 6A-B).** TNF-α increased significantly in both frequency and MFI, while IFN-γ, GM-CSF, and IL-17 were each significantly increased in either frequency or MFI **(Fig. 6C-F)**. These findings indicate that inhibition of WNT signaling partially restores CD4^+^ T-cell activation and inflammatory cytokine expression in the presence of Tregs.

To determine whether soluble factors contributed to this effect, we exposed CFSE-labeled CD4^+^ T cells to supernatants from Treg or Th0 cultures in the presence or absence of mDKK-1. Cells cultured alone served as controls, and the gating strategy is shown in **Fig. S8**. In control conditions, CD4^+^ T cells displayed baseline proliferation and CD25 expression that were not significantly affected by mDKK-1 treatment **(Fig. 7A)**. In contrast, in the presence of Treg-conditioned supernatant, mDKK-1 treatment significantly increased both CD4^+^ T cell proliferation and CD25 expression **(Fig. 7B).** No comparable effects were observed with Th0-conditioned supernatant, where neither proliferation nor CD25 expression was significantly altered by mDKK-1 **(Fig. 7C)**. To further assess cytokine production, we measured secreted IFN-γ, GM-CSF, TNF-α, and IL-10 by ELISA (**Fig. S9**). WNT inhibition increased pro-inflammatory cytokines in Treg-conditioned cultures, whereas IL-10 remained unchanged. Similar but more modest effects were observed in Th0-conditioned cultures.

**Figure 7.**
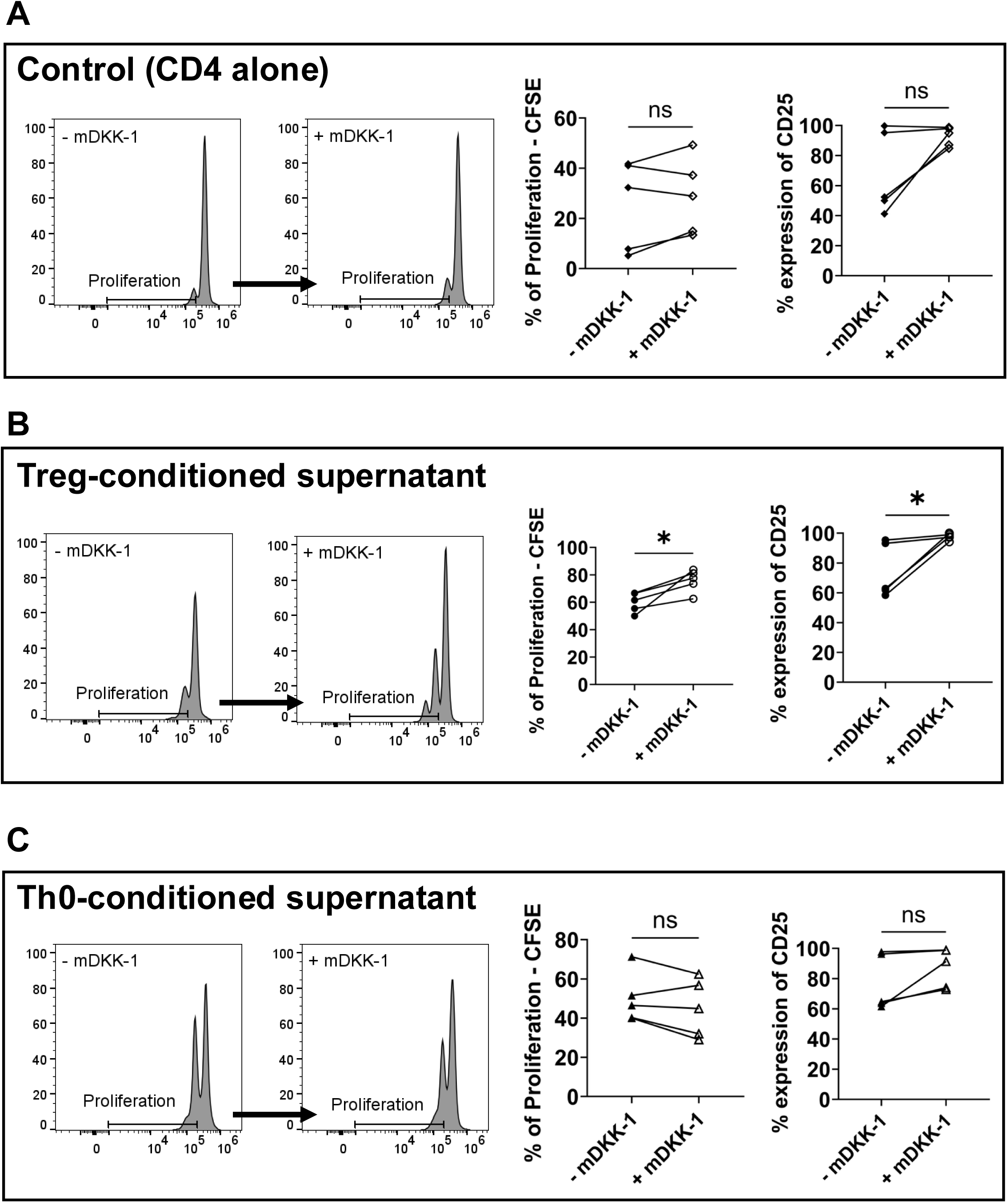
Soluble Treg-derived factors regulate CD4^+^ T cell proliferation and activation in a WNT-dependent manner. CFSE dilution and CD25 expression were analyzed in CD4^+^ T cells stimulated with anti-CD3 and anti-CD28 antibodies and cultured under the indicated conditions in the presence or absence of mDKK-1. (**A**) CD4^+^ T cells cultured alone, (**B**) CD4^+^ T cells cultured with Treg-conditioned supernatant, and (**C**) CD4^+^T cells cultured with Th0-conditioned supernatant. For each condition, representative CFSE histograms are shown together with quantification of proliferation and CD25 expression. Data represent mean ± SEM from five independent experiments (n = 5). Statistical significance was determined using paired t-tests (\**p* < 0.05).

Taken together, these supernatant-transfer experiments indicate that soluble Treg-derived factors contribute to WNT-dependent regulation of CD4^+^ T cell responses, with the clearest effects observed on proliferation and CD25 expression.

## 4. Discussion and Conclusion

Tregs are central to immune homeostasis, limiting excessive effector T cell activation and preventing inflammatory pathology. Although multiple suppressive mechanisms have been described, including cytokine secretion, metabolic disruption, and inhibitory receptor signaling^1, 30, 31^, additional molecular pathways likely contribute to Treg function. Here, we identify canonical WNT signaling as a previously underappreciated component of the Treg regulatory program. Tregs preferentially expressed and secreted several canonical WNT ligands, including WNT2B, WNT3, WNT7B, and WNT10B, whereas activated CD4^+^ T cells expressed Frizzled receptors capable of sensing these signals. Moreover, pharmacological inhibition of canonical WNT signaling partially reversed Treg-mediated suppression of CD4^+^ T cell activation, proliferation, and inflammatory cytokine production. Together, these findings support the existence of a paracrine signaling axis in which Treg-derived WNT ligands likely contribute to the regulation of effector CD4^+^ T cell responses.

WNT ligands have previously been detected in several immune cell populations, including macrophages, dendritic cells, and stromal cells, where they participate in regulating immune activation, tissue repair, and inflammatory responses^14, 32–36^. However, the specific production of WNT ligands by CD4^+^ T cell subsets has remained largely unexplored. Our data demonstrate that Tregs display a distinct WNT expression profile compared with naïve and effector CD4^+^ T cells. Both transcriptional and protein analyses revealed enrichment of canonical WNT ligands in Tregs, suggesting that WNT production may represent an additional component of the Treg suppressive program.

Although WNT7B and WNT10B were detected intracellularly in nTregs, secreted forms of these proteins were not reliably quantified by ELISA. Unlike WNT2B and WNT3, which were detectable in conditioned media under our experimental conditions, the recovery of certain WNT ligands in soluble assays can be limited by the intrinsic biochemical properties of WNT proteins. WNT proteins are lipid-modified secreted glycoproteins that undergo palmitoylation by the acyltransferase PORCN and are trafficked through a Wntless (WLS)-dependent secretion pathway, processes that confer significant hydrophobicity and often promote membrane association or vesicle-mediated transport^37–40^. These properties complicate the detection of specific WNT ligands in conventional immunoassays and may indicate that some WNT molecules preferentially act through short-range mechanisms, including membrane tethering or extracellular vesicle-mediated signaling. Such localized signaling could allow Tregs to deliver spatially restricted regulatory cues to neighboring T cells, thereby limiting effector responses without broadly suppressing the surrounding immune environment. Together, these observations expand the repertoire of molecules associated with Treg-mediated immune regulation and suggest that WNT production may contribute to the establishment of immunoregulatory microenvironments.

Although Th0 cells produced detectable levels of certain WNT proteins, their secretion was consistently lower than that observed in Tregs, indicating that regulatory T cells represent a prominent source of these ligands within CD4^+^ T cell populations. Notably, *WNT2B* and *WNT3* transcripts were also elevated in Th0 cells, whereas ELISA demonstrated greater accumulation of these proteins in Treg-conditioned supernatants. This suggests that regulation of these ligands may occur not only at the transcriptional level but also at the level of protein processing, trafficking, or secretion. More broadly, these findings raise the possibility that activated CD4^+^ T cells themselves may also engage WNT-dependent regulatory circuits. This interpretation is consistent with our observation that mDKK-1 enhanced activation and pro-inflammatory cytokine expression even in CD4^+^ T cells cultured alone, and with prior reports that activated T cells can secrete WNT ligands under specific stimulatory conditions^15^. Together, these data suggest that canonical WNT signaling may function both as a Treg-derived paracrine mechanism and as an autocrine or activation-associated regulatory pathway within conventional CD4^+^ T cells.

Canonical WNT/β-catenin signaling has previously been implicated in multiple aspects of CD4^+^ T cell biology, including cell survival, differentiation, and memory formation^25, 41^. Our findings extend these observations by demonstrating that WNT signaling also modulates the magnitude of CD4^+^ T cell activation and inflammatory cytokine production. Inhibition of canonical WNT signaling using mDKK-1 enhanced the expression of activation markers and increased production of pro-inflammatory cytokines such as IFN-γ, TNF-α, GM-CSF, and IL-17 in activated CD4^+^ T cells. These results suggest that canonical WNT signaling functions as a regulatory mechanism that restrains excessive CD4^+^ T cell activation. Notably, the effects of WNT inhibition were more pronounced in the presence of Tregs, where blockade of WNT signaling partially reversed Treg-mediated suppression of effector T cell responses. These findings support a model in which Treg-derived WNT ligands likely contribute to the suppression of inflammatory T cell responses through canonical WNT signaling in neighboring CD4^+^ T cells.

The selective effects of WNT signaling observed in our study further highlight the specificity of this regulatory pathway. While inhibition of WNT signaling increased the production of several pro-inflammatory cytokines, IL-10 secretion remained largely unchanged. This observation suggests that canonical WNT/β-catenin signaling does not broadly suppress all aspects of CD4^+^ T cell function but instead selectively restrains inflammatory effector pathways. IL-10 expression in CD4^+^ T cells is largely regulated by transcription factors such as STAT3, c-Maf, and IRF4 downstream of cytokine receptor and TCR signaling^42–45^, pathways that are largely independent of canonical WNT signaling. This regulatory architecture may explain why inhibition of WNT signaling enhanced pro-inflammatory cytokine production while leaving IL-10 levels unaffected, allowing Tregs to limit excessive effector responses while preserving anti-inflammatory mechanisms that support immune resolution.

Functional assays using Treg-conditioned supernatants further reinforced the importance of soluble WNT molecules in mediating these effects. Exposure of CD4^+^ T cells to Treg-derived supernatants altered proliferation and activation in a WNT-dependent manner, as pharmacological inhibition of WNT signaling restored both proliferative capacity and CD25 expression. In contrast, Th0-derived supernatants produced limited effects on proliferation and activation, consistent with the absence of significant changes in CFSE dilution and CD25 expression under these conditions. Together, these findings support a model in which soluble WNT molecules produced by Tregs act in a paracrine manner to restrain effector CD4^+^ T cell responses.

Despite these insights, several limitations of the current study should be considered. First, inhibition of WNT signaling was achieved using the pharmacological antagonist mDKK-1, which blocks canonical WNT signaling through inhibition of LRP5/6 co-receptors. Although this approach is widely used to disrupt WNT/β-catenin signaling, it may also influence additional signaling pathways or interact with other regulatory networks involved in immune responses^46, 47^. Future studies using genetic approaches, such as conditional deletion of specific WNT ligands in Tregs, would provide greater specificity and allow direct assessment of the contribution of individual WNT molecules to Treg-mediated suppression.

Second, although multiple WNT ligands were identified as enriched in Tregs, the relative contribution of each ligand to the observed immunomodulatory effects remains unclear. Canonical WNT ligands can exhibit overlapping or context-dependent functions, and it is possible that several WNT proteins act cooperatively to shape T cell responses. Dissecting the individual roles of specific WNT ligands through targeted knockdown or conditional knockout approaches will therefore be important for refining our understanding of how WNT signaling contributes to Treg function.

Finally, our study was performed primarily using *in vitro* systems, which allowed controlled investigation of WNT-mediated interactions between Tregs and CD4^+^ T cells. Future work extending these findings to *in vivo* models of immune activation or inflammatory disease will be essential to determine the physiological relevance of Treg-derived WNT signaling in regulating immune responses within complex tissue environments.

In conclusion, our study identifies Tregs as a previously underrecognized source of canonical WNT ligands and demonstrates that these molecules contribute to the suppression of effector CD4^+^ T cell responses (**Fig. 8**). By limiting proliferation, activation, and pro-inflammatory cytokine production, Treg-derived WNT signaling represents an additional mechanism through which regulatory T cells maintain immune homeostasis. These findings expand current understanding of Treg-mediated immune regulation and identify WNT signaling as a previously underappreciated component of the Treg suppressive program, providing a conceptual framework for targeting this pathway to modulate inflammatory T cell responses.

**Figure 8.**
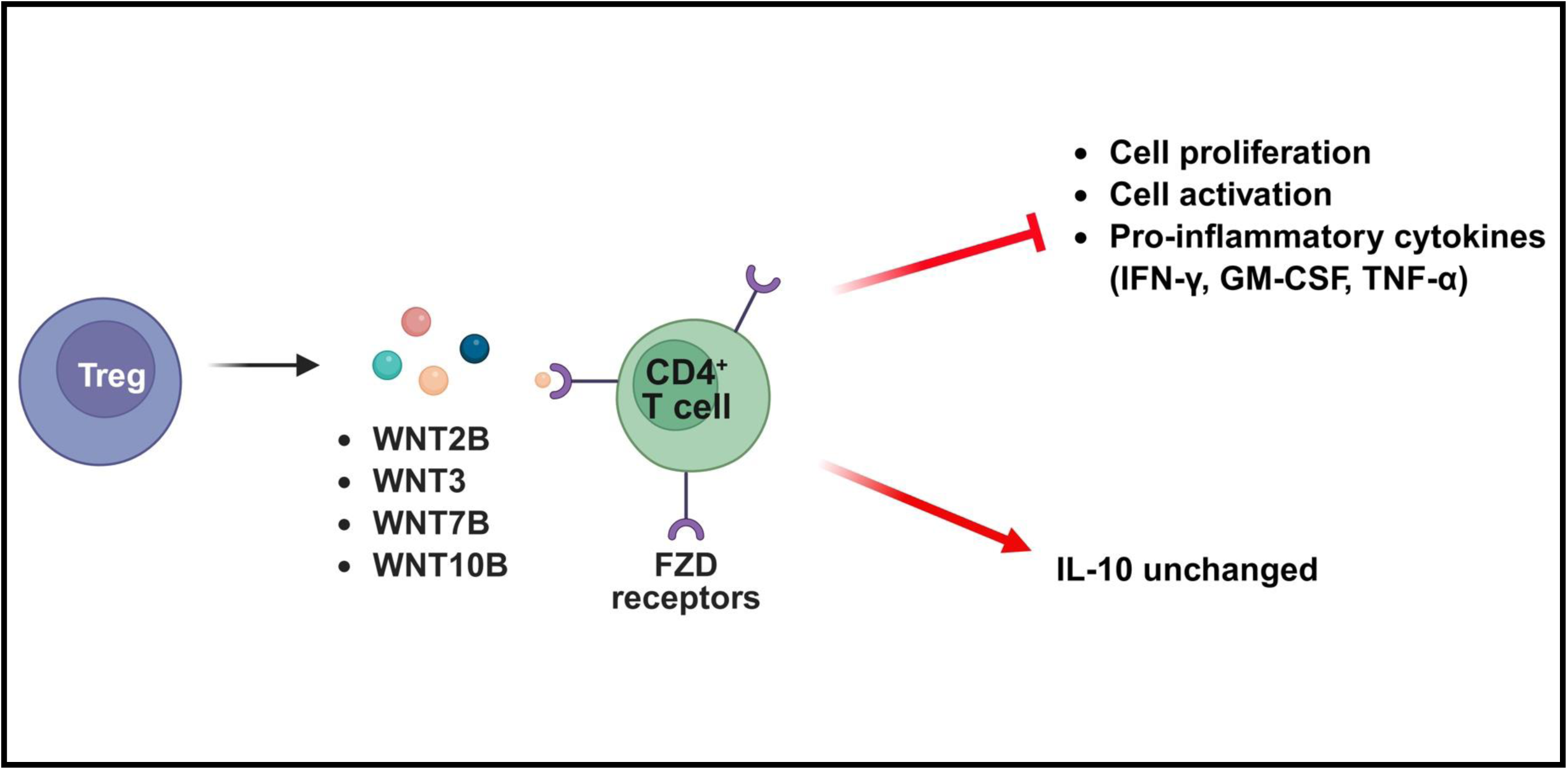
Proposed model of WNT-mediated regulation of CD4^+^ T cell responses by Tregs. Regulatory T cells (Tregs) express and secrete WNT ligands, whereas conventional CD4^+^ T cells express Frizzled (FZD) receptors capable of sensing these signals. Treg-derived WNT molecules are proposed to limit effector CD4^+^ T cell responses, resulting in reduced proliferation, decreased activation, and diminished pro-inflammatory cytokine expression, while IL-10 remains unchanged. Together, these findings support a model in which Treg-derived WNT signaling contributes to the regulation of effector CD4^+^ T cell responses.

## Acknowledgements

We thank the Research Flow Cytometry Facility in the Division of Rheumatology at Cincinnati Children’s Hospital Medical Center for providing access to instrumentation for flow cytometry data acquisition.

## Fundings

This work was supported by grants from the National Institutes of Health Grant R35GM146890 (II) and R03NS144870 (II).

## Conflict of Interest

The authors declare no conflict of interest.

## Author contributions

I. I. and K.S.P. conceptualized the study and designed the experiments. K.S.P. performed the majority of the experiments, including mouse tissue harvest, cell isolation, cell differentiation, *ex vivo* treatment administration, and flow cytometry, and conducted data analysis and visualization. M. N. performed the Frizzled (FZD) receptor expression experiments and conducted flow cytometry for these studies. K.S.P. and I.I. drafted and revised the manuscript. All authors read, edited, and approved the final version of the manuscript.

## Data availability

All data generated or analyzed during this study are included in the manuscript and its supplementary information files.

## Notes

### Competing Interest Statement

The authors have declared no competing interest.

