## Supplemental Information for "Canonical WNT Ligands Produced by Regulatory T Cells Restrain Effector CD4^+^ T Cell Responses"

#### SUPPLEMENTAL FIGURE LEGENDS

**Figure S1. Gating strategy for identification of naïve $CD4^+$ T cells.** Representative flow cytometry dot plots illustrate the sequential gating strategy used to identify naïve $CD4^+$ T cells from splenocytes. Debris and doublets were excluded, and viable lymphocytes were selected prior to gating on $CD4^+$ T cells. Naïve $CD4^+$ T cells were defined as $CD4^+CD44^{lo}CD62L^{hi}CD25^-FOXP3^-$ and are highlighted by a red box.

**Figure S2. Gating strategy for identification of differentiated regulatory T cells (Tregs).** Representative flow cytometry dot plots illustrate the sequential gating strategy used to identify differentiated regulatory T cells (Tregs). Following exclusion of debris and doublets and selection of viable lymphocytes, $CD4^+$ T cells were gated. Tregs were defined as $CD4^+CD25^+FOXP3^+$ and are highlighted by a red box.

**Figure S3. WNT gene expression profile in Tregs compared with naïve $CD4^+$ T cells.** Heatmap showing the relative expression of WNT genes in Tregs compared with naïve $CD4^+$ T cells (NCT) (n = 4). Gene expression was normalized to the average of the housekeeping genes *B2m* and *Actb*. Values are presented as log2 fold change; red indicates increased expression and green indicates decreased expression in Tregs relative to NCT. The color scale ranges from +5 to -3.

**Table 1. Differential expression of WNT genes in Tregs and naïve $CD4^+$ T cells.** WNT gene expression was assessed by qPCR in Tregs and naïve $CD4^+$ T cells, with normalization to the average of the housekeeping genes *B2m* and *Actb*. Fold-regulation values and corresponding p-values are shown for each gene. Genes significantly upregulated in Tregs ($p < 0.05$), including *WNT2B, WNT3,* and *WNT10B*, are highlighted in red. *WNT7B*, which showed a trend toward significance ($p = 0.07$), is highlighted in pink. *WNT9B* served as a negative control ($p = 0.18$) and is highlighted in blue.

**Figure S4. Gating strategy for identification of naïve CD4$^{+}$ T cells and natural regulatory T cells (nTregs).** Representative flow cytometry plots illustrate the sequential gating strategy used to identify naïve CD4$^{+}$ T cells and natural regulatory T cells (nTregs) from splenocytes. Cells were gated to exclude debris (**A**), select singlets (**B**), and identify viable lymphocytes (**C**), followed by gating on CD4$^{+}$ T cells (**D**). Naïve CD4$^{+}$ T cells were defined as CD44$^{lo}$CD62L$^{hi}$CD25$^{-}$FOXP3$^{-}$(**E-F**), whereas nTregs were defined as CD4$^{+}$CD25$^{+}$FOXP3$^{+}$ cells (**G**). This gating strategy was used for analysis of WNT protein expression shown in Fig. 2.

**Figure S5. Gating strategy for identification of differentiated Th0 cells.** Representative flow cytometry dot plots illustrate the sequential gating strategy used to identify Th0 cells. Debris and doublets were excluded, and viable lymphocytes were selected prior to gating on CD4$^{+}$ T cells. Th0 cells were defined as CD4$^{+}$CD25$^{+}$FOXP3$^{-}$ and are highlighted by a red box.

**Figure S6. Gating strategy for identification of conventional CD4$^{+}$ T cells**. Representative flow cytometry dot plots illustrate the sequential gating strategy used to identify conventional $CD4^+$ T cells. Debris and doublets were excluded, and viable lymphocytes were selected prior to gating on $CD4^+$ T cells. Regulatory T cells ($CD4^+CD25^+FOXP3^+$, blue gate) were excluded, and the remaining conventional $CD4^+$ T cells ($CD4^+CD25^-FOXP3^-$, red gate) were used for downstream analyses.

**Figure S7. Representative flow cytometry plots of $CD4^+$ T cell activation and cytokine expression in co-culture with Tregs ± mDKK-1**. Representative flow cytometry plots illustrate the analysis of conventional $CD4^+$ T cells under four culture conditions: (**A**) $CD4^+$ T cells cultured alone; (**B**) $CD4^+$ T cells cultured alone with mDKK-1; (**C**) $CD4^+$ T cells co-cultured 1:1 with Tregs; and (**D**) $CD4^+$ T cells co-cultured 1:1 with Tregs in the presence of mDKK-1. The top row shows activation markers CD69 and CD25, the middle row shows TNF-α and IFN-γ, and the bottom row shows GM-CSF and IL-17. Plots are representative of four independent experiments (n = 4).

**Figure S8. Gating strategy for analysis of $CD4^+$ T cell proliferation and activation**. Representative flow cytometry dot plots illustrate the sequential gating strategy used for analysis of $CD4^+$ T cell proliferation and activation. Debris and doublets were excluded, and viable lymphocytes were selected prior to gating on $CD4^+$ T cells. The gated $CD4^+$ T cell population was used for downstream analysis of CFSE dilution and CD25 expression.

**Figure S9. Cytokine production by $CD4^+$ T cells exposed to Treg- or Th0-conditioned supernatants ± mDKK-1.** $CD4^+$ T cells were stimulated with anti-CD3 and anti-CD28 antibodies and cultured with conditioned media from differentiated Tregs or

Th0 cells in the presence or absence of mDKK-1. Cytokine production was quantified by ELISA. Shown are levels of (**A**) IFN-γ, (**B**) GM-CSF, (**C**) TNF-α, and (**D**) IL-10 (pg/mL). Data represent mean ± SEM from five independent experiments ($n = 5$). Statistical significance was determined using paired t-tests (*$p < 0.05$, **$p < 0.01$).

### Figure S1

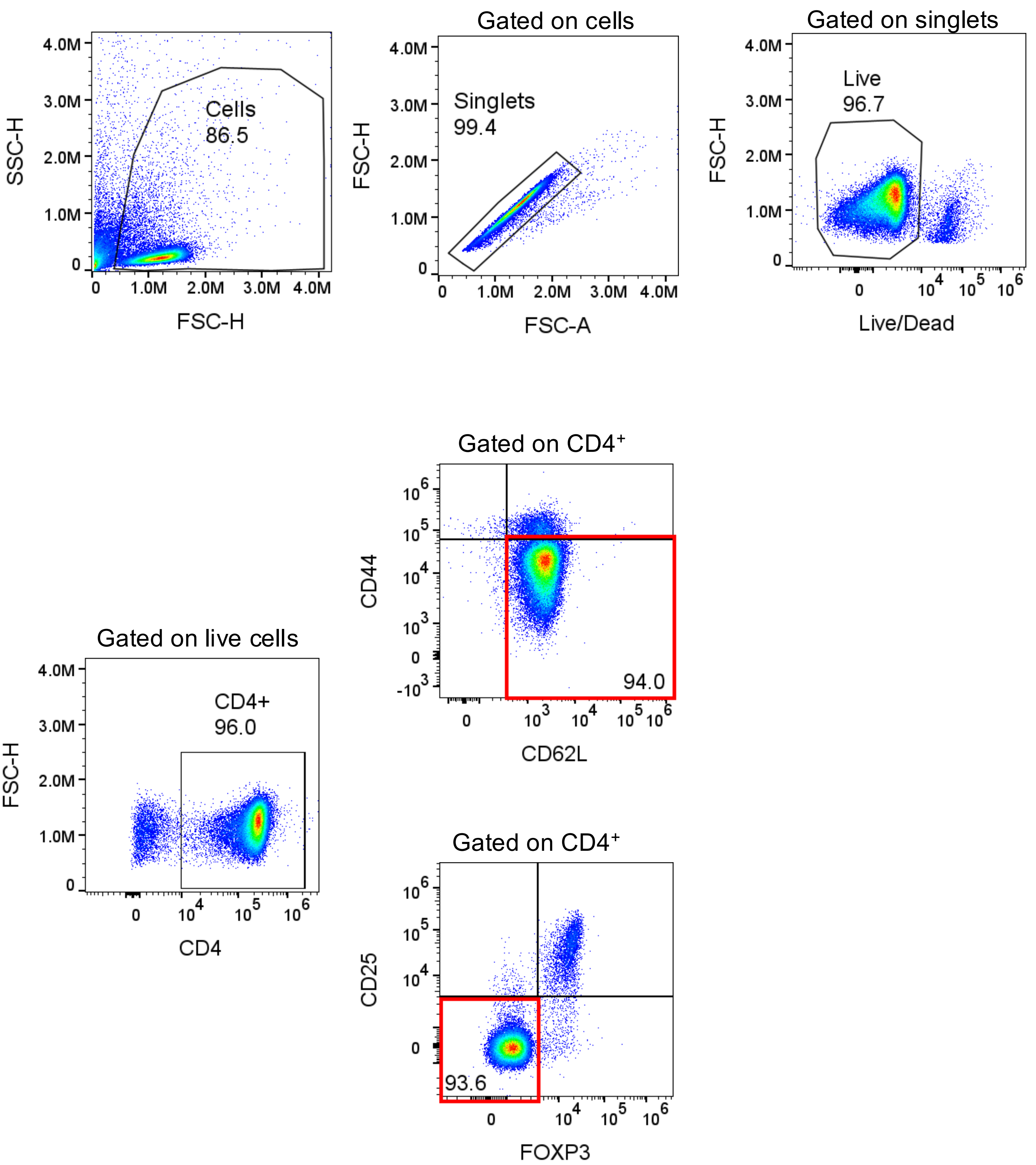
Gated on cells
Gated on singlets
Cells
86.5
SSC-H
FSC-H
Singlets
99.4
FSC-A
Live
96.7
Live/Dead
Gated on live cells
CD4+
96.0
CD4
Gated on CD4⁺
94.0
CD44
CD62L
93.6
CD25
FOXP3

#### Figure S2

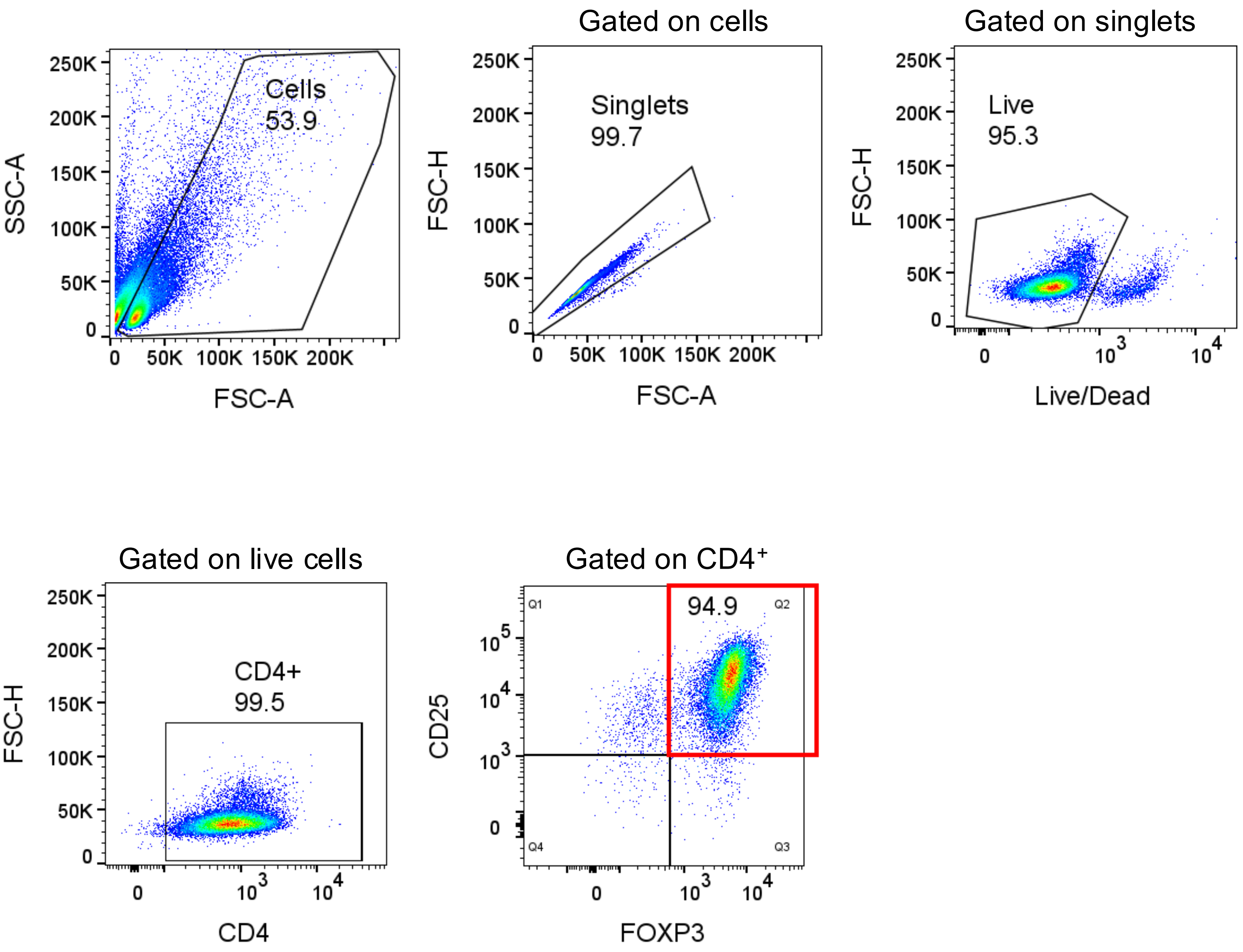

Cells
53.9
SSC-A
FSC-A
250K
200K
150K
100K
50K
0
50K
100K
150K
200K
Gated on cells
Singlets
99.7
FSC-H
FSC-A
Gated on singlets
Live
95.3
FSC-H
Live/Dead
$10^3$
$10^4$
Gated on live cells
CD4+
99.5
FSC-H
CD4
Gated on CD4+
94.9
Q1
Q2
Q3
Q4
CD25
FOXP3
$10^5$

**Figure S3**

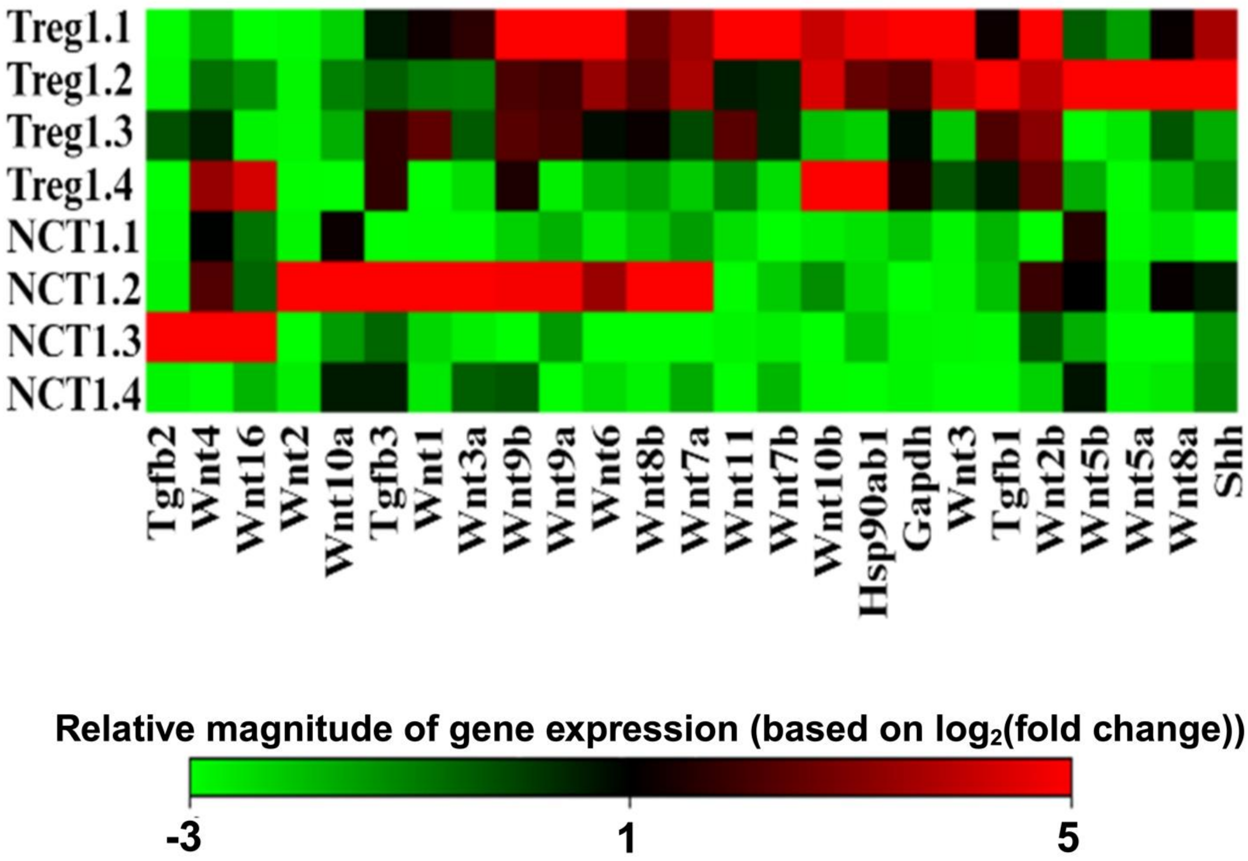

Treg1.1
Treg1.2
Treg1.3
Treg1.4
NCT1.1
NCT1.2
NCT1.3
NCT1.4
Tgfb2
Wnt4
Wnt16
Wnt2
Wnt10a
Tgfb3
Wnt1
Wnt3a
Wnt9b
Wnt9a
Wnt6
Wnt8b
Wnt7a
Wnt11
Wnt7b
Wnt10b
Hsp90ab1
Gapdh
Wnt3
Tgfb1
Wnt2b
Wnt5b
Wnt5a
Wnt8a
Shh
Relative magnitude of gene expression (based on $log_2$(fold change))
-3
1
5

**Table 1**

| Gene | Fold regulation | p-value |
|---|---|---|
| WNT1 | 1.82 | 0.766901 |
| WNT3A | 2.05 | 0.913717 |
| WNT6 | 3.53 | 0.204013 |
| WNT8B | 2.52 | 0.412157 |
| WNT10B | 9.58 | 0.019540 |
| Tgfb1 | 5.38 | 0.003532 |
| WNT7A | 2.01 | 0.537743 |
| WNT9A | 1.93 | 0.458324 |
| WNT11 | 32.35 | 0.011616 |
| WNT2B | 3.28 | 0.009353 |
| WNT5A | 5.20 | 0.260121 |
| WNT7B | 6.22 | 0.079197 |
| WNT9B | 4.21 | 0.182107 |
| WNT3 | 24.51 | 0.038476 |
| WNT8A | 4.96 | 0.172399 |

#### Figure S4

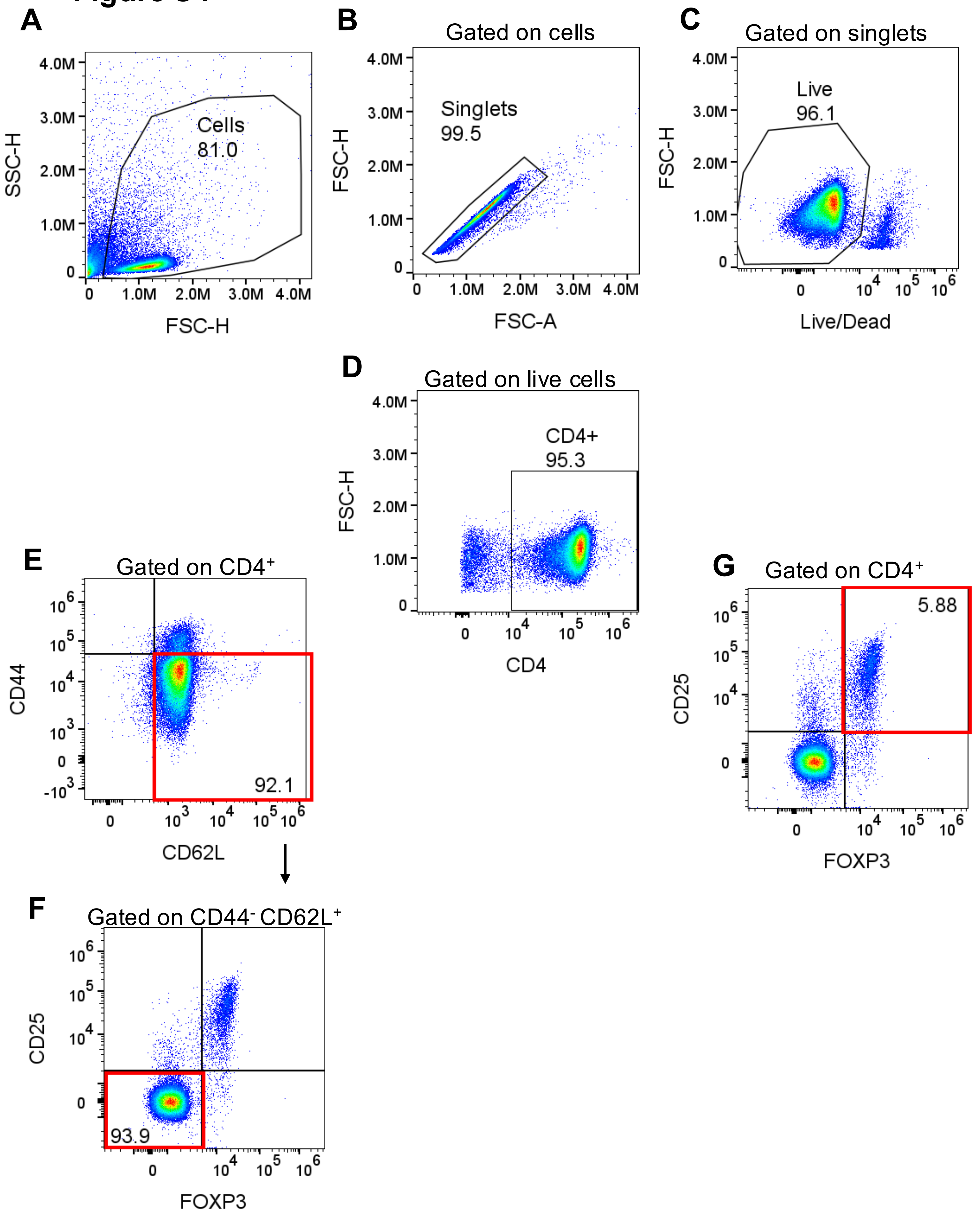
A
Cells
81.0
SSC-H
FSC-H
B
Gated on cells
Singlets
99.5
FSC-H
FSC-A
C
Gated on singlets
Live
96.1
FSC-H
Live/Dead
D
Gated on live cells
CD4+
95.3
FSC-H
CD4
E
Gated on CD4+
CD44
CD62L
92.1
F
Gated on CD44- CD62L+
CD25
FOXP3
93.9
G
Gated on CD4+
5.88
CD25
FOXP3

### Figure S5

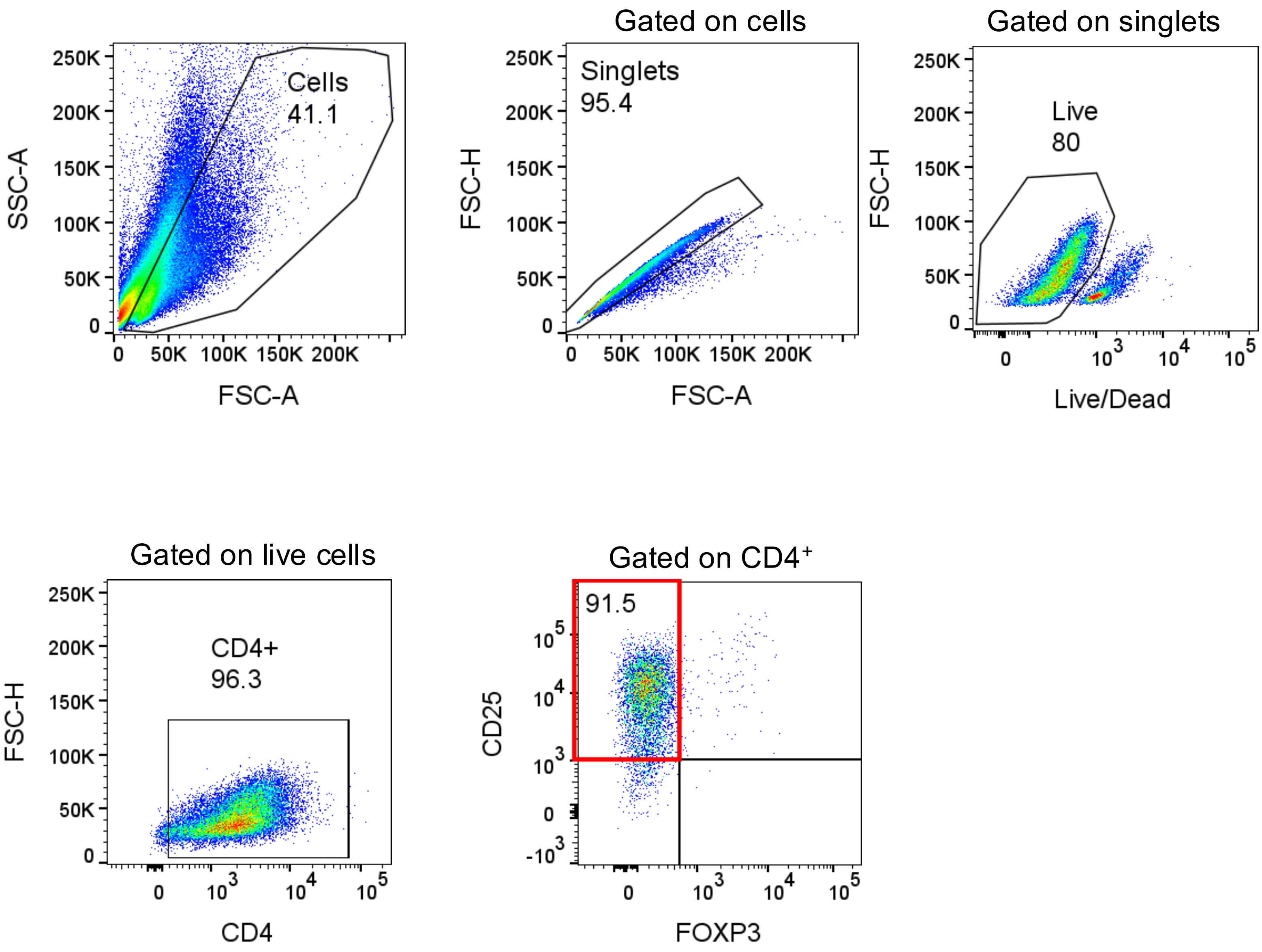

Gated on cells
Gated on singlets
Cells
41.1
Singlets
95.4
Live
80
SSC-A
FSC-A
FSC-H
Live/Dead
Gated on live cells
Gated on CD4⁺
CD4+
96.3
91.5
CD4
FOXP3
CD25

#### Figure S6

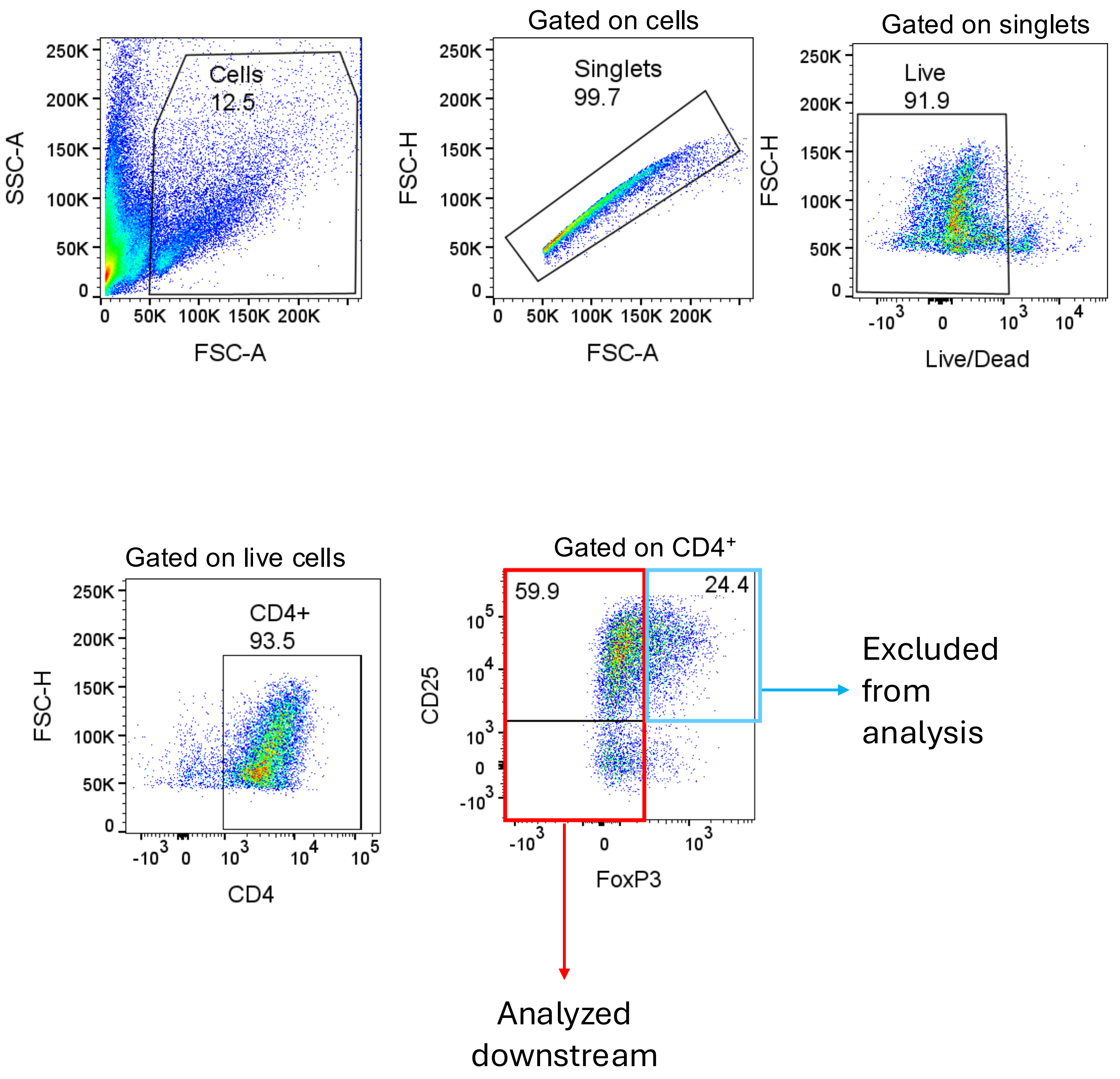
Gated on cells
Gated on singlets
Cells
12.5
Singlets
99.7
Live
91.9
SSC-A
FSC-A
FSC-H
Live/Dead
Gated on live cells
CD4+
93.5
CD4
Gated on CD4⁺
59.9
24.4
CD25
FoxP3
Excluded from analysis
Analyzed downstream

#### Figure S7

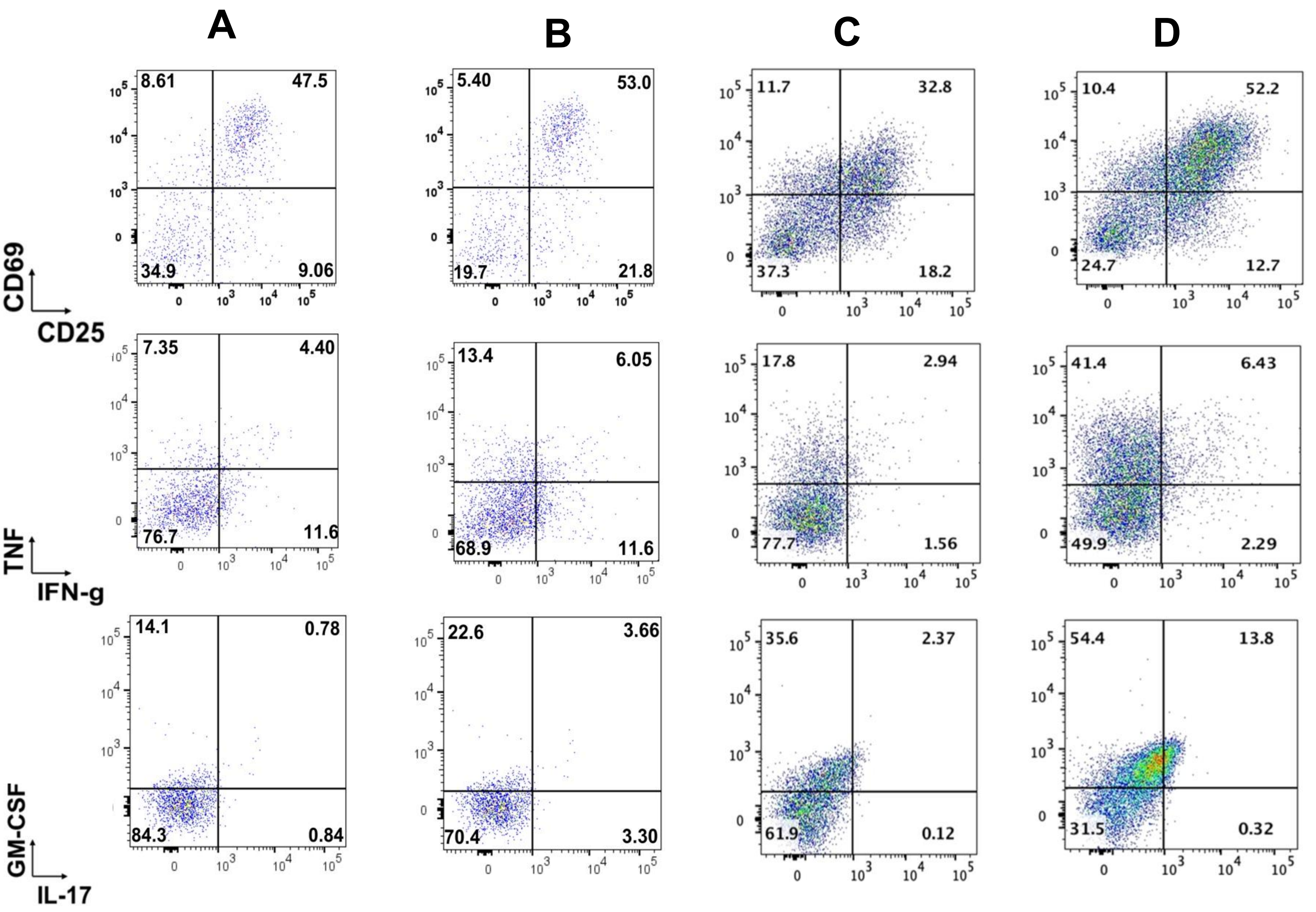


| | | | | |
|---|---|---|---|---|
| $CD4^+$ T cells | + | + | + | + |
| Tregs | - | - | + | + |
| mDKK-1 | - | + | - | + |

### Figure S8

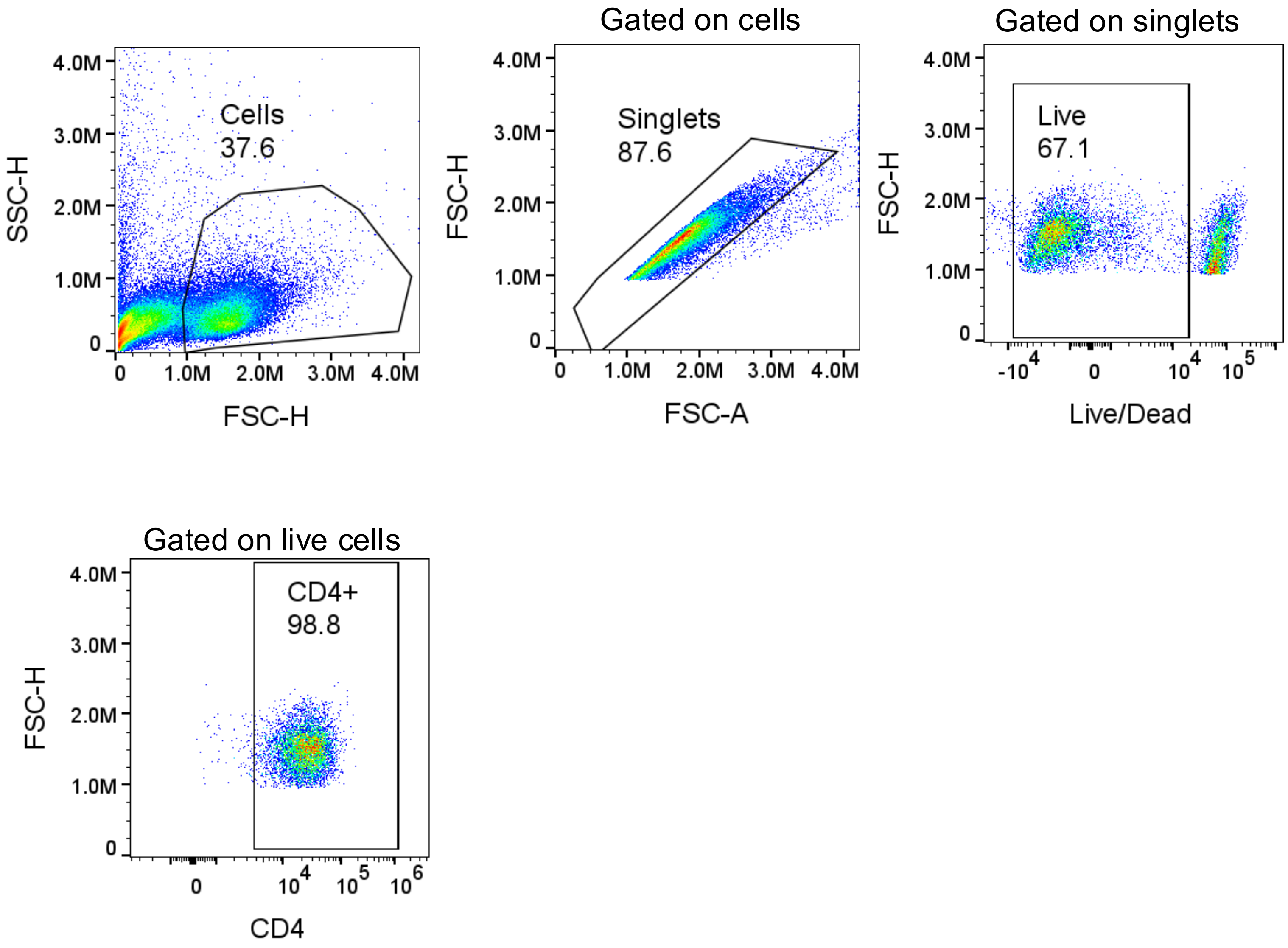

Cells
37.6
SSC-H
FSC-H
Gated on cells
Singlets
87.6
FSC-H
FSC-A
Gated on singlets
Live
67.1
FSC-H
Live/Dead
Gated on live cells
CD4+
98.8
FSC-H
CD4

#### Figure S9

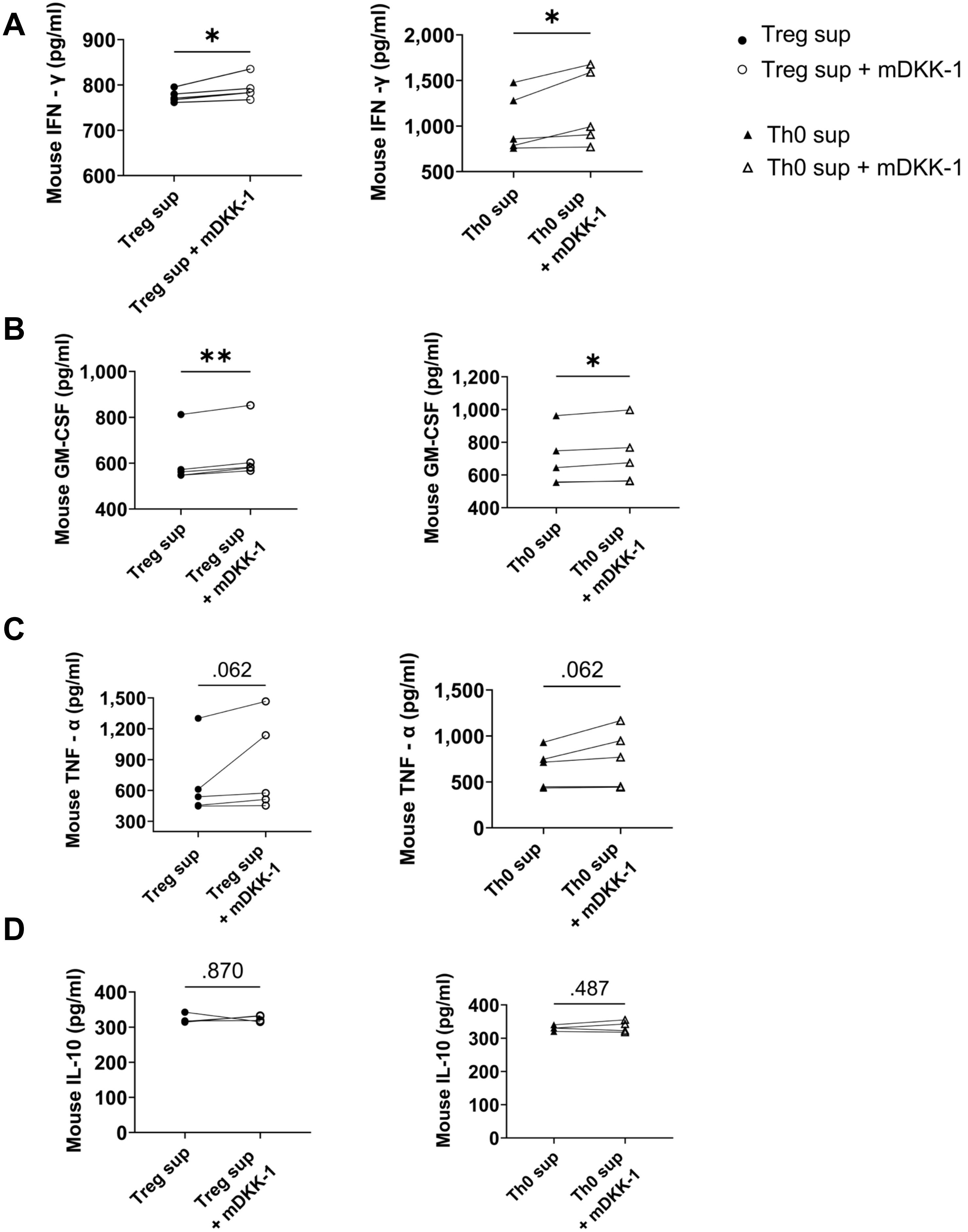
A
Mouse IFN - γ (pg/ml)
Treg sup
Treg sup + mDKK-1
Mouse IFN -γ (pg/ml)
Th0 sup
Th0 sup + mDKK-1
Treg sup
Treg sup + mDKK-1
Th0 sup
Th0 sup + mDKK-1
B
Mouse GM-CSF (pg/ml)
C
Mouse TNF - α (pg/ml)
.062
D
Mouse IL-10 (pg/ml)
.870
.487
